# Improving DASH adherence after midlife promotes better late-life cognition and brain health: the SG70 Study

**DOI:** 10.64898/2026.09.23.753372

**Authors:** Eric Kwun Kei Ng, Jun S. Lai, Fang Ji, Yi Liang Benjamin Goh, Phoebe Si Qi Chia, Wen Liang Loh, Joice Sin Yi Tham, Sheng Fong, Woon-Puay Koh, Juan Helen Zhou

## Abstract

Healthier dietary patterns reduce cognitive impairment risks, but whether diet improvement after mid-life can benefit cognition and brain health, and how brain measures mediate diet–cognition associations, remain unclear, as few studies had long-term diet records alongside detailed neuroimaging and cognitive measures. We assessed adherence to the Dietary Approaches to Stop Hypertension (DASH) in 892 SG70 participants, nested within the Singapore Chinese Health Study, between 1993–2025. Adherence during midlife (44–55 years) and early elderhood (61–73 years) were related to seven cognitive domains, brain morphometry, white matter hyperintensities (WMH), and free-water markers measured in late life (68–82 years). Higher adherence at both mid-life stages was associated with better verbal memory. Higher mid-life adherence was associated with greater white matter volume in association tracts, whereas higher early-elderhood adherence was associated with lower free water (frontal and occipital) and lower WMH (basal ganglia and anterior periventricular). Moreover, white matter volume and free water mediated the diet-verbal memory associations at their respective midlife stages. Importantly, participants with improved mid-to-late life diet trajectories showed better late-life verbal memory and brain health than those with persistently low adherence. Overall, ensuring mid-life diet quality and improving post-midlife diet may promote cognitive resilience through structural reserve and neurovascular health.

## INTRODUCTION

Amid rapid global ageing, the gap between healthspan and lifespan is widening ^1^ Cognitive decline and neurodegenerative diseases such as dementia represent major threats to healthspan and exacerbate clinical and socioeconomic burdens. Modifiable risk factors offer opportunities for intervention ^2^, with diet emerging as a safe and promising target ^3^. Healthy dietary patterns such as the Dietary Approaches to Stop Hypertension (DASH) ^4^ show protective effects with broad applicability in both Western and Asian cohorts ^56^. However, a key challenge remains in understanding the long-term impact of diet quality in earlier adulthood on later-life cognition and brain health. In particular, midlife represents a period of significant yet modifiable biological transitions that steer ageing trajectories ^7–9^. Examining dietary history and longitudinal dietary trajectories enable the delineation of stage-dependent effects ^10^, suggest directional influence ^11^, and reveals actionable targets after midlife ^12^.

Evidence on such long-term influence of diet on neurocognitive ageing remains incomplete ^13^. Many past longitudinal studies focus on short to intermediate follow-up periods that exclude midlife, or use a single diet measure instead of examining its time-varying impact ^5,13,14^. Studies with long-term follow-ups or multiple diet records ^9,11,15,16^ are often devoted to investigating dietary benefits against cognitive impairment and dementia without neuroimaging, restricting insights into the neurophysiological underpinnings. Moreover, cognitive vulnerability varies across domains ^17^, but domain-specific cognition was not always assessed ^5,15,18^, making it uncertain if healthy dietary patterns can selectively benefit late-life cognition ^19,20^, particularly in Asian populations ^5^. Finally, investigation of the impact of dietary changes in later life on brain and cognitive health remains inconclusive ^21^. A comprehensive life-course framework integrating diet, neuroimaging, and cognition is essential to tackle these outstanding questions.

Related, the neurobiology linking dietary patterns to brain ageing is also poorly understood ^3,6,18^, as evidenced by the heterogenous findings from magnetic resonance imaging (MRI) studies ^14,22^. On one hand, shorter follow-up periods in existing work prohibit conclusions on the stage-dependent and long-term dietary effects ^13,14^. On the other hand, while macroscopic measures such as brain morphometry and total white matter hyperintensity (WMH) burden ^14,23^ remain indispensable for studying structural integrity and cerebrovascular disease, respectively, in the context of dietary research, recent methodological advances have not been incorporated. Specifically, it is unknown if free water, a novel diffusion MRI marker sensitive to neurovascular inflammation and subtle microstructural changes ^16,24,25^, and the spatial distribution of WMH, which may reflect distinct vascular or non-vascular origins ^26^, can deepen our understanding of dietary influences on the brain. Importantly, linking diet-sensitive brain phenotypes and cognitive outcomes through mediation can help identify neural substrates that relate diet to late-life cognitive resilience, yet such analyses are few and have yielded inconclusive results so far ^27,28^.

To address these gaps, we analysed data from a cohort of 892 community-dwelling Chinese older adults living in Singapore in the SG70 study. Participants had repeated assessments of diet quality spanning up to 28 years across adulthood (ages 44–55, 61–73, and 68–82 years), alongside comprehensive cognitive testing and multimodal brain MRI in late life (ages 68–82 years). We hypothesised that greater adherence to the DASH pattern during midlife and early elderhood would be associated with superior late-life cognitive performance. With respect to brain health, we further hypothesised that better historical diet quality would be associated with more favourable brain morphometry and lower level of cerebrovascular burden and neurovascular inflammation. Such diet-associated brain phenotypes would mediate the relationship between earlier-life diet quality and late-life cognitive outcomes. Finally, we expected that participants would display different DASH adherence patterns from mid to late life, and those who had a more favourable trajectory would show better cognitive and/or brain health in late life.

## RESULTS

The SG70 study is nested within the population-based Singapore Chinese Health Study, which recruited community-dwelling Chinese adults in Singapore between 1993 and 1998, and followed them up over multiple visits, including follow-ups in 2014–2016 and 2021–2025. ^29^. At this latest follow-up, 1,165 participants aged 68-82 years underwent multidimensional assessments of ageing outcomes, forming the SG70 subcohort (**Fig. 1A**) ^30^. For the present analyses, we included 892 SG70 participants (489 women) with largely complete multimodal MRI and domain-specific neuropsychological data (**Table 1**). They also had complete dietary assessments in midlife (ages 44–55 years) at recruitment, early elderhood (ages 61–73 years) and late-life (ages 68-82 years) at follow-up, thus enabling a life-course analysis of DASH adherence in relation to late-life brain and cognitive phenotypes (**Fig. 1B**) ^31^.

**Figure 1.**
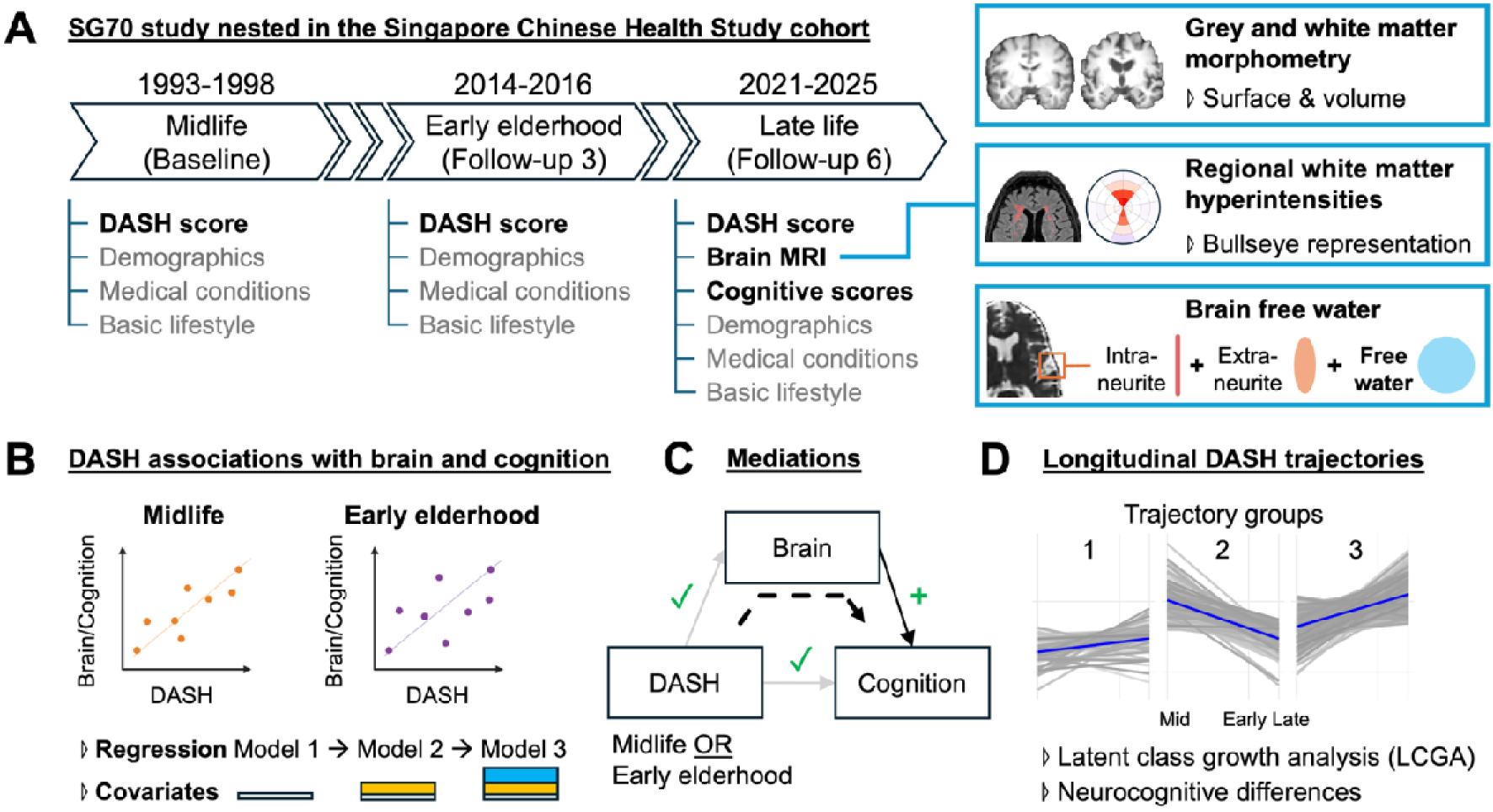
Schematic illustration of the study design. **A)** Singaporean Chinese adults were enrolled in the Singapore Chinese Health Study (SCHS) during midlife (44-55 years) and followed longitudinally across multiple time points. Dietary intake, used to derive DASH adherence, together with demographic, lifestyle, and health variables was collected at selected visits. At the most recent follow-up in late life (the SG70 study; follow-up 6; 68-82 years), participants additionally underwent multimodal brain magnetic resonance imaging (MRI) and comprehensive neuropsychological assessment across seven cognitive domains. **B)** To examine associations between life-stage diet quality and late-life neurocognitive outcomes, cognitive domain scores and brain measures, including morphometry, cerebrovascular burden, and free water, were related to DASH adherence in midlife (baseline) and early elderhood (follow-up 3) using multiple regression models with progressive covariate adjustment (Model 1-3; see **Methods**). Cross-sectional analyses with both DASH and neurocognitive measures in late life were also conducted (not shown). **C)** To investigate potential diet-brain-cognition (dotted black arrows), neurocognitive measures that showed significant associations with diet in earlier life stages (paths with green ticks) and brain-cognition associations (paths with green plus signs) were subject to mediation analysis. **D)** Finally, to assess the impact of dietary changes after midlife, longitudinal DASH trajectories across all time points were derived using latent class growth analysis. Neurocognitive measures implicated in the mediation analyses were then compared across selected trajectory groups, with a priori focus on those with improved diets after midlife.

**Table 1.** Characteristics of the SG70 participants across adulthood.

|  | Midlife (baseline) | Early elderhood (follow-up 3) | Late life (follow-up 6) |
| --- | --- | --- | --- |
|  | N = 892 <sup>a</sup> |  |  |
| Sex (all visits) |  |  |  |
| M |  | 403 (45.2%) |  |
| F |  | 489 (54.8%) |  |
| Dialect group (all visits) |  |  |  |
| Cantonese |  | 439 (49.2%) |  |
| Hokkien |  | 453 (50.8%) |  |
| Education (all visits) |  |  |  |
| No |  | 48 (5.4%) |  |
| Primary |  | 267 (29.9%) |  |
| Secondary |  | 414 (46.4%) |  |
| Tertiary |  | 163 (18.3%) |  |
| Age, years (Mean ± SD) | 47.5 (± 2.23) | 67.2 (± 2.39) | 74.3 (± 2.63) |
| DASH score (Mean ± SD) | 20.8 (± 4.60) | 21.1 (± 4.67) | 21.7 (± 4.63) |
| Weekly participation in strenuous sports |  |  |  |
| No | 741 (83.1%) |  |  |
| 0.5-1hr | 77 (8.6%) |  |  |
| 2-3hs | 51 (5.7%) |  |  |
| 4+hr | 23 (2.6%) |  |  |
| Weekly participation in vigorous work |  |  |  |
| No | 805 (90.2%) |  |  |
| 5-3hrs | 53 (5.9%) |  |  |
| 4-6hrs | 13 (1.5%) |  |  |
| 7+hr | 21 (2.4%) |  |  |
| Amount of moderate activity per week |  |  |  |
| No | 681 (76.3%) |  |  |
| 0.5-3hrs | 159 (17.8%) |  |  |
| 4+hr | 52 (5.8%) |  |  |
| Amount of physical activity per week |  |  |  |
| No |  | 70 (7.9%) |  |
| 0.5-4hrs |  | 156 (17.5%) |  |
| 4+hr |  | 666 (74.7%) |  |
| Alcohol consumption |  |  |  |
| Never/Occasional | 776 (87%) | 783 (87.8%) | 797 (89.3%) |
| Weekly | 102 (11.4%) | 84 (9.4%) | 77 (8.6%) |
| Daily | 14 (1.6%) | 25 (2.8%) | 18 (2.0%) |
| Smoking history |  |  |  |
| Never | 760 (85.2%) | 737 (82.6%) | 742 (83.2%) |
| Ex | 57 (6.4%) | 110 (12.3%) | 118 (13.2%) |
| Current | 75 (8.4%) | 45 (5.0%) | 32 (3.6%) |
| Amount of sleep per day, hours |  |  |  |
| 7hrs | 336 (37.7%) | 270 (30.3%) | 272 (30.5%) |
| 6hrs | 209 (23.4%) | 254 (28.5%) | 194 (21.7%) |
| 5hrs | 52 (5.8%) | 156 (17.5%) | 27 (3.0%) |
| 8hrs | 250 (28%) | 169 (18.9%) | 305 (34.2%) |
| 9hrs | 45 (5%) | 43 (4.8%) | 77 (8.6%) |
| Missing |  |  | 17 (1.9%) |
| Body mass index |  |  |  |
| Underweight [$< 18.5 \text{ kg/m}^2$] | 58 (6.5%) | 390 (43.7%) | 52 (5.8%) |
| Normal [$18.5 \text{ to } 22.9 \text{ kg/m}^2$] | 446 (50.0%) | 57 (6.4%) | 314 (35.2%) |
| Overweight [$23.0 \text{ to } 27.4 \text{ kg/m}^2$] | 330 (37.0%) | 358 (40.1%) | 400 (44.8%) |
| Obese [$\geq 27.5 \text{ kg/m}^2$] | 58 (6.5%) | 82 (9.2%) | 125 (14.0%) |
| Hypertension | 99 (11.1%) | 444 (49.8%) | 534 (59.9%) |
| Heart and artery diseases | 8 (0.9%) | 51 (5.7%) | 61 (6.8%) |
| Stroke | 2 (0.2%) | 18 (2.0%) | 34 (3.8%) |
| Diabetes | 18 (2.0%) | 145 (16.3%) | 221 (24.8%) |
| Cancer | 11 (1.2%) | 57 (6.4%) | 95 (10.7%) |
| Cognition [Min, Max] <sup>b</sup> |  |  |  |
| Verbal memory |  |  | [23.4, 72.7] |
| Visual memory |  |  | [30.2, 74.7] |
| Missing |  |  | 3 |
| Attention |  |  | [21.7, 78.0] |
| Visuospatial ability |  |  | [25.2, 85.8] |
| Executive functioning |  |  | [31.1, 79.9] |
| Missing |  |  | 28 |
| Processing speed |  |  | [21.8, 78.1] |
| Missing |  |  | 15 |
| Language |  |  | [28.0, 68.4] |
| Missing |  |  | 1 |
| MRI structural T1 |  |  | 892 |
| MRI T2-FLAIR |  |  | 889 |
| MRI multi-shell diffusion |  |  | 881 |

Note: <sup>a</sup> Participants who completed dietary questionnaires, neuropsychological assessments, and provided good
quality MRI data; <sup>b</sup> All cognitive T scores were normalised to have M = 50, SD = 10.

### Midlife and early elderhood diet quality influenced future cognitive performance in late life in a domain-specific manner

Extending prior findings with global cognition ^11,29^, we first examined the relationships between DASH adherence at each life stage and performance across seven cognitive domains in late life ^30,32,33^, namely processing speed, attention, visuospatial ability, executive function, visual memory, verbal memory, and language ability. We found that higher DASH adherence in midlife (**Fig. 2A**), early elderhood (**Fig. 2B**), and late life was consistently associated with better late-life verbal memory after multiple comparison correction (**Supplementary Table 1**). Notably, the association with midlife DASH adherence remained significant when all life-stage DASH measures were modelled jointly ^31^, indicating an enduring contribution of earlier-life diet to late-life cognitive function (**Supplementary Table 3**).

**Figure 2.**
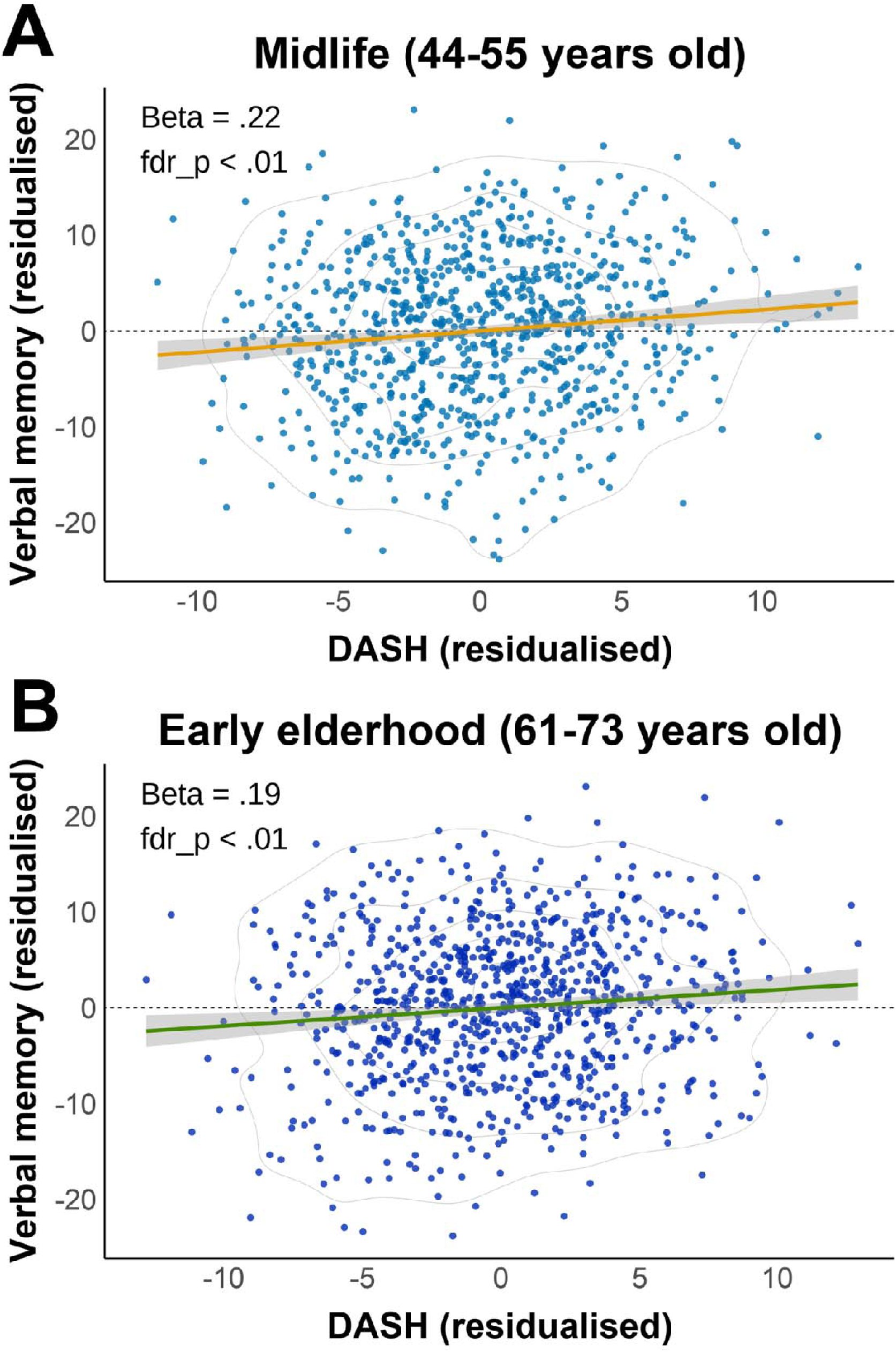
Better life-stage diet quality was associated with better verbal memory in late life. The scatter plots show the association between verbal memory in late life and DASH adherence in midlife (44-55 years; **A**) and early elderhood (61-73 years; **B**), respectively. The cross-sectional association between DASH adherence and verbal memory in late life (68 – 82 years) was also statistically significant (see main text). Notably, when DASH adherence across all life stages was modelled jointly, higher midlife DASH adherence remained independently associated with better late-life verbal memory (see main text). Variables in the plots are residualised for demographic, health, and lifestyle covariates and adjusted for multiple comparisons across seven cognitive domains. The grey shading represents the 95% confidence interval of the estimated linear fit. Contour lines indicate data distribution at the 5^th^, 25^th^, 50^th^, 75^th^, and 95^th^ percentiles.

### Better diet quality in midlife was associated with larger white matter volume in late life

We next examined the associations between DASH adherence at each life stage and late-life brain morphometry. In white matter, higher midlife DASH adherence was associated with greater white matter volume in the corpus callosum, inferior frontal and parietotemporal regions (**Fig. 3A**, cluster-level familywise error correction p < .05). Anatomically, these clusters encompassed major association tracts, including the superior longitudinal fasciculus, arcuate fasciculus, and parietal aslant tracts, which are implicated in language-related processing ^34^ (**Supplementary Fig. 1**).

**Figure 3.**
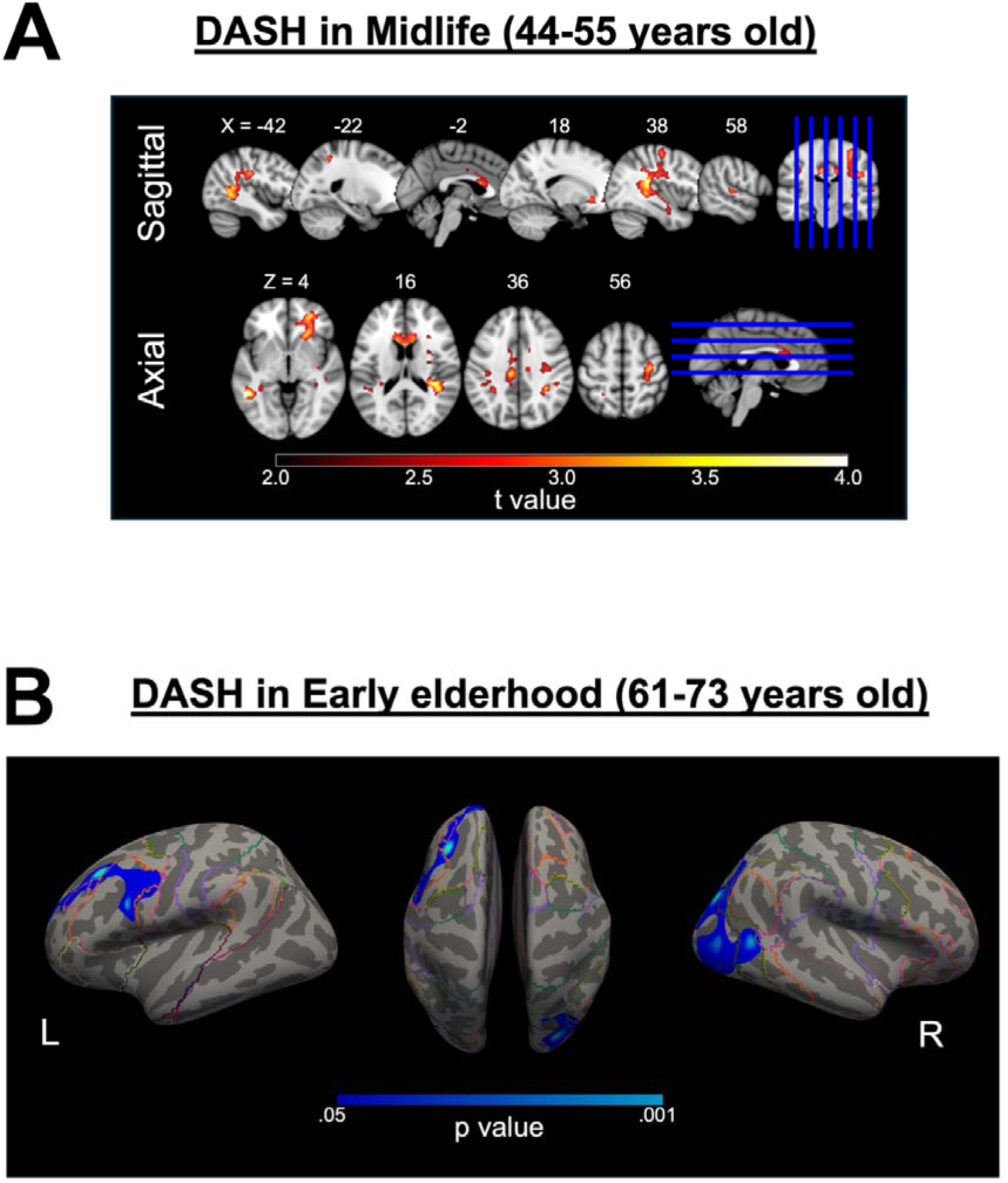
Higher diet quality in midlife was associated with better white matter morphometry in late life, while higher diet quality in early elderhood was associated with lower cortical grey matter free water in late life. **A)** Distributed white matter volume clusters in the corpus callosum, inferior frontal, and parietotemporal regions were significantly associated with higher DASH adherence in midlife. **B)** Lower free water in the left prefrontal and right occipital cortices was significantly associated with higher DASH adherence in early elderhood. Results are after adjustment for key past and recent demographic, self-reported health, and lifestyle covariates (see Methods). Numbers above each brain slice in (A) indicate MNI X (sagittal) or Z (axial) coordinates.

Findings from grey matter measures were more modest. Higher DASH adherence in early elderhood was associated with greater cortical thickness in the left precuneus and increased volume in somatomotor regions, whereas higher late-life DASH adherence was associated with larger bilateral inferior frontal volumes (**Supplementary Fig. 2**).

### Better diet quality in early elderhood was associated with lower brain grey matter free water in late life

To investigate whether midlife diet quality was related to late life brain microstructure, we related life-stage DASH adherence to voxel-wise/vertex-wise free water measures in cortical grey matter, subcortical grey matter, and skeletonised white matter ^35,36^. We found that higher DASH adherence in early elderhood was associated with lower cortical grey matter free water in left frontal and right occipital regions (**Fig. 3B**; cluster-level familywise error correction p < .05), spanning default mode, control, attention, and visual networks ^37^ (**Supplementary Fig. 3**). Similar inverse associations were also observed cross-sectionally between late-life DASH adherence and late-life cortical free water, with a more widespread cortical distribution (**Supplementary Fig. 4**). No significant associations were detected in subcortical grey matter or white matter tracts.

### Better diet quality in early elderhood was associated with lower regional white matter hyperintensity burden in late life

We then examined the associations between DASH adherence at different life stages and regional white matter hyperintensity (WMH) volume in late life, a robust marker of cerebrovascular pathology. We focused on seven spatial components of regional WMHs (**Supplementary Fig. 5**), reflecting distinct aetiological patterns as implicated in prior studies ^26^. Higher DASH adherence in early elderhood was associated with lower WMH burden in the deep basal ganglia and to a lesser extent anterior periventricular regions (**Fig. 4**; **Supplementary results – White matter hyperintensities**). These findings suggest that higher diet quality in earlier adulthood may help mitigate cerebrovascular burden in late life.

**Figure 4.**
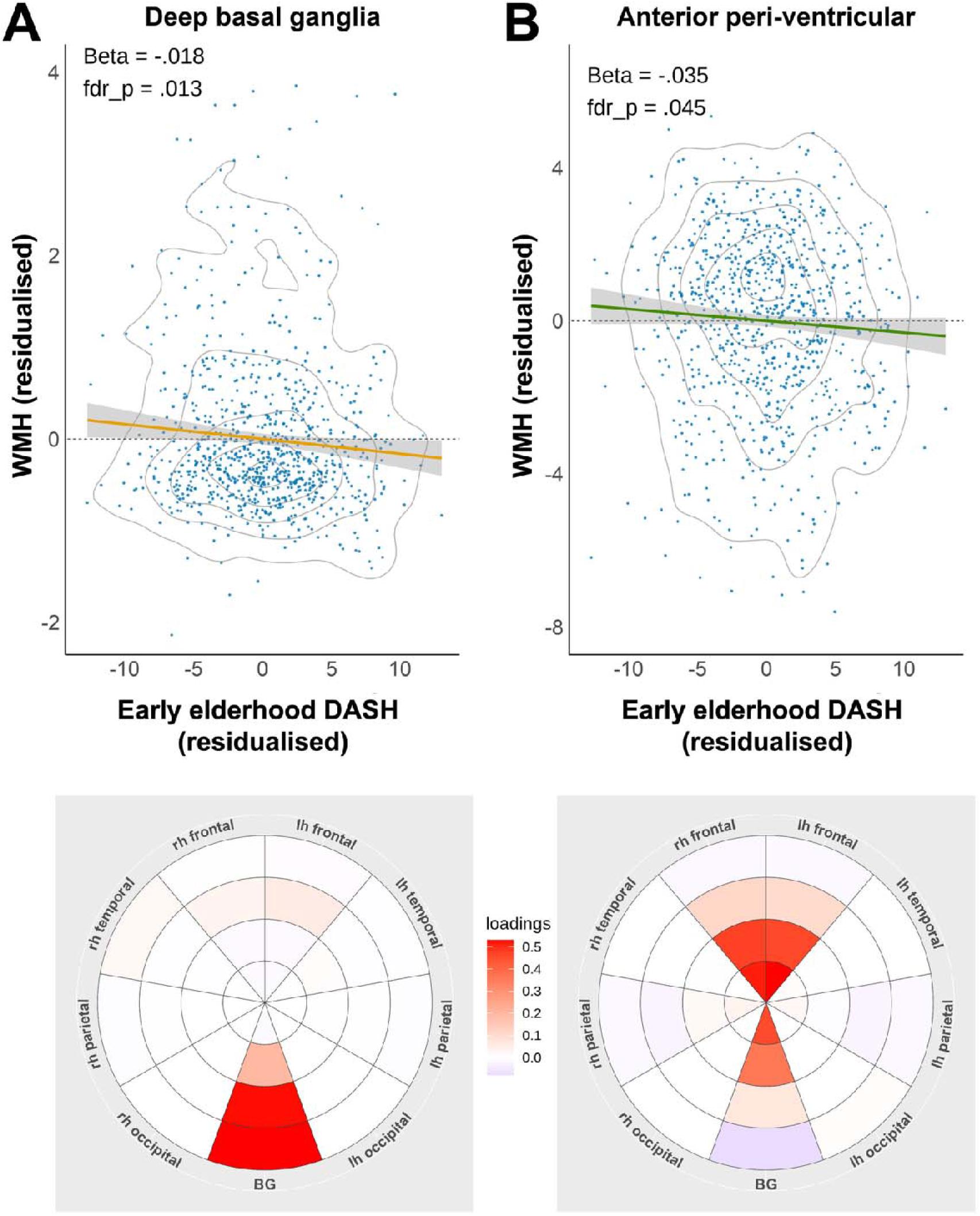
Higher diet quality in early elderhood was associated with lower regional white matter hyperintensities (WMH) at late life. **(A)** Scatter plot (top) shows the associations between DASH adherence in early elderhood and WMH factor scores for the white matter around deep basal ganglia, whose spatial distribution is depicted in the bullseye representation (bottom). **(B)** Same summary for the anterior periventricular WMH. All analyses were adjusted for basic demographic covariates and intracranial volume. Shades in the scatter plots represent the 95% confidence intervals of the fitted regression lines. Contour lines indicate data distribution at the 5^th^, 25^th^, 50^th^, 75^th^, and 95^th^ percentiles. For the bullseye representations, concentric rings represent four equidistant zones between the ventricles and the grey matter-white matter boundary, transversing peri-ventricular, deep, and juxta-cortical depths of the white matter. These zones are further divided by lobes (frontal, temporal, parietal, occipital, and basal ganglia) and hemispheres (bilateral except basal ganglia). Right hemisphere (rh) is displayed on the left.

### Brain phenotypes mediated the relationship between mid-life diet and late-life verbal memory

To establish relevance to cognition, we examined the associations between diet-related brain measures and verbal memory, all measured at late life. Higher white matter volume and lower cortical free water were associated with better verbal memory (**Supplementary Table 11**), whereas regional WMH burden and grey matter morphometry were not significantly associated with verbal memory. These findings implicate white matter structure and brain free water as candidate neurobiological correlates linking earlier-life diet quality to late-life cognitive health.

Moreover, mediation analyses (**Fig. 1C**, **Methods – mediation analysis**) revealed that white matter volume partially mediated the association between midlife DASH adherence and late-life verbal memory (indirect effect = 0.027, bootstrapped 95% CI = 0.008–0.052; 12.3% mediated; **Fig. 5A**). Similarly, cortical free water partially mediated the association between higher DASH adherence in early elderhood and late-life verbal memory (indirect effect = 0.023, bootstrapped 95% CI = 0.007–0.044; 12.5% mediated; **Fig. 5B**). Together, these findings suggest a mediating role of brain health in the protective effects of historical diet on late life cognition.

**Figure 5.**
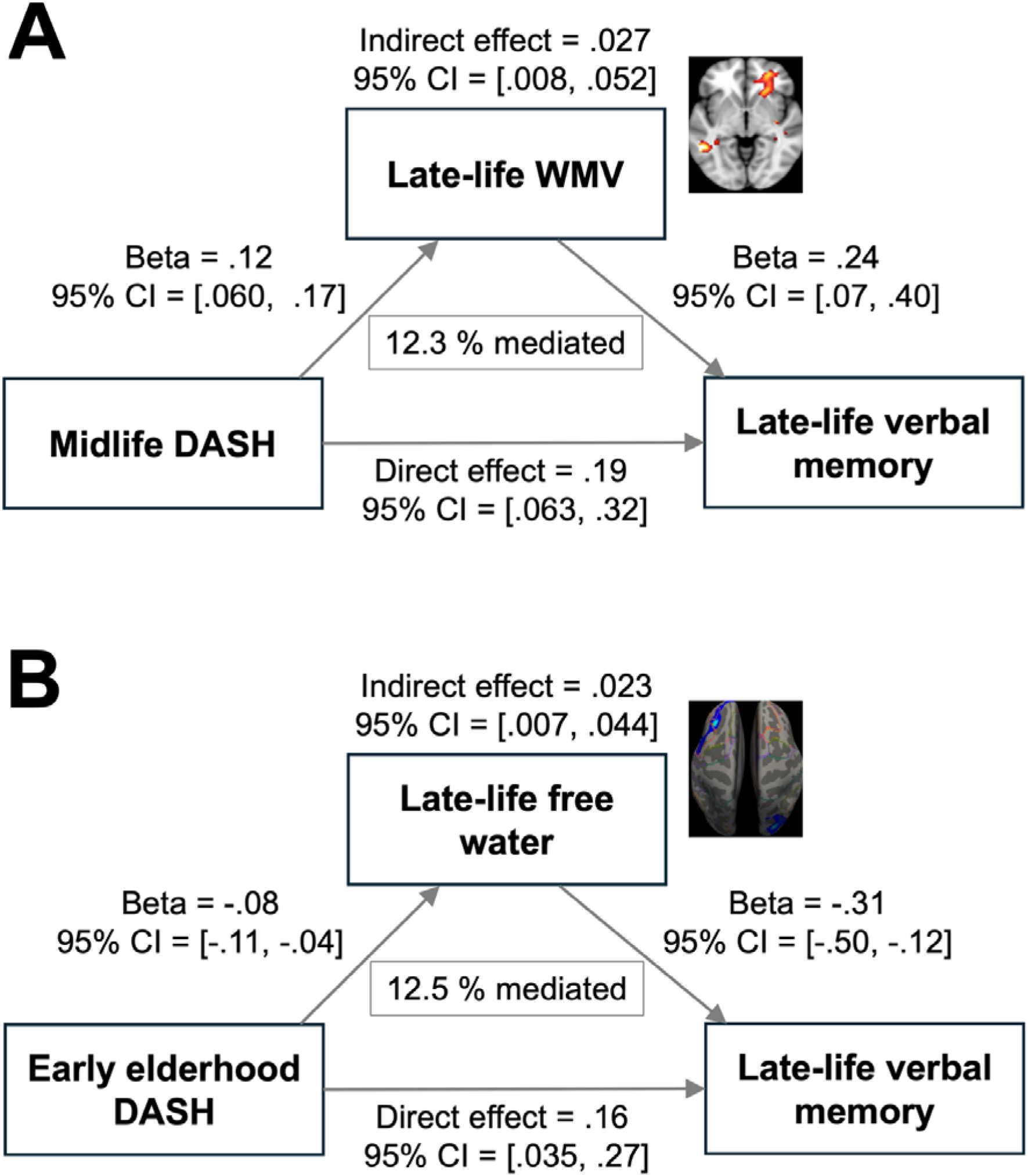
Late-life brain white matter structural health and grey matter neurovascular health mediated the associations between better life-stage diet quality and late-life verbal memory. **A)** Greater regional white matter volume partially mediated the association between higher mid-life DASH adherence and better late-life verbal memory (12.3% mediated). **B)** Lower free water in these regions partially mediated the association between higher early-elderhood DASH adherence and better late-life verbal memory (12.5% mediated). Averaged white matter volumes and free water from respective significant clusters were used in the mediation analyses. Path coefficients are shown with bootstrapped 95% confidence intervals, and the proportion mediated is calculated based on unstandardised coefficients. Estimates are multiplied by 100 for more compact numerical presentation only.

### Diet improvement after midlife still conferred benefits to brain and cognitive health at late life

Finally, as understanding dietary patterns in midlife and subsequent improvements may help define actionable windows for intervention ^12^, we identified six DASH trajectory groups using latent class growth analysis across the three life stages (**Fig. 1D; Methods – DASH trajectory derivation and analyses; Supplementary Table 12**). These trajectories ranged from consistently low to consistently high DASH adherence, with two groups showing improvement after midlife despite low or moderate diet quality at the beginning (low-increasing and average-increasing; **Fig. 6A, Supplementary Table 13**).

**Figure 6.**
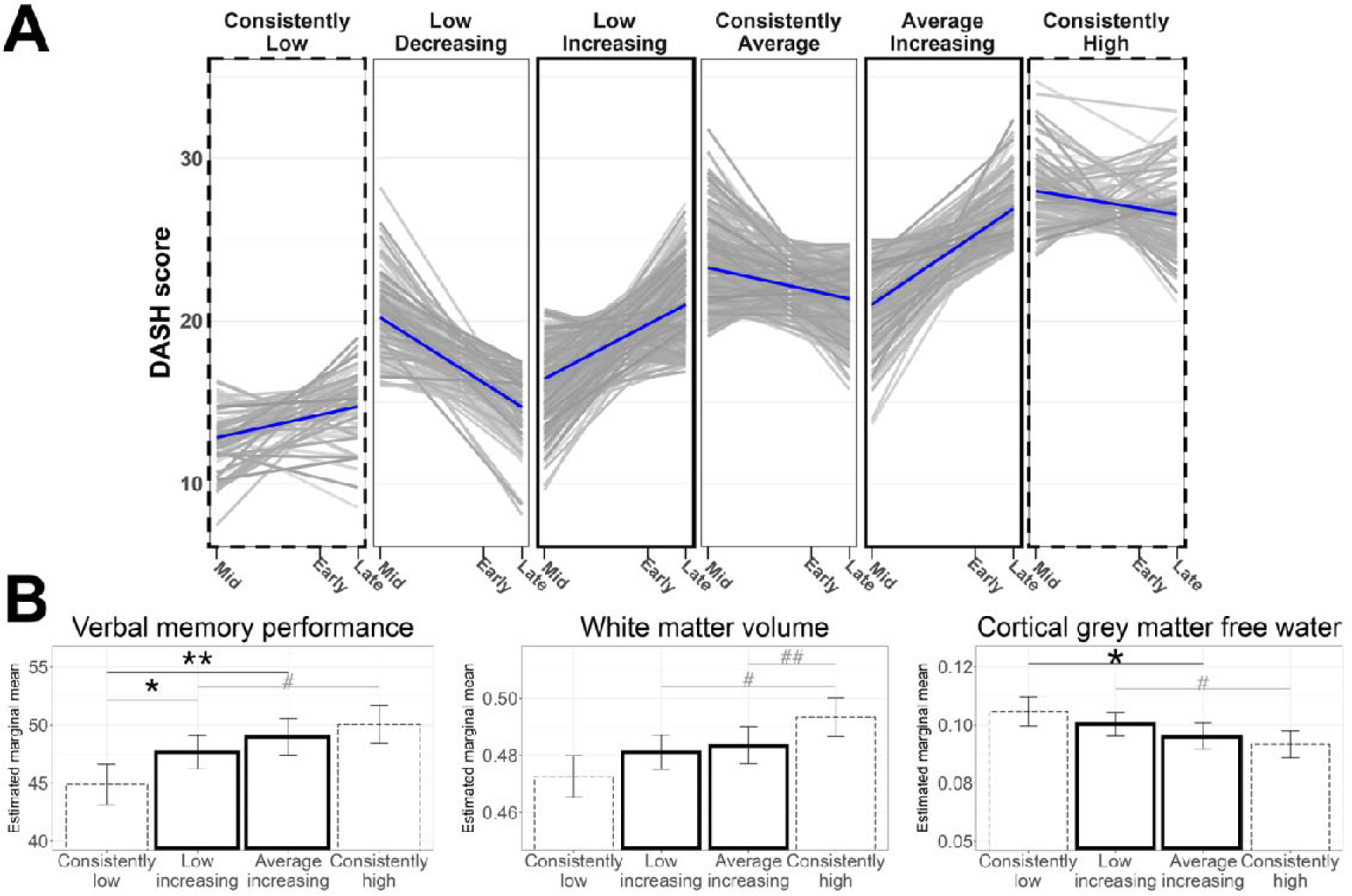
Improving diet quality after midlife was associated with better late-life cognitive and brain health. **A)** Latent class growth analysis identified six dietary trajectory groups across adulthood (blue lines represent estimated group trajectories; grey lines represent individual trajectories), with two groups showing significant increase (Low increasing, Average increasing) and one showing significant decrease (Low decreasing) in diet quality after midlife (slope ps < .001). The two ‘increasing’ groups (thick black panels) were subsequently compared to the two anchor groups (dotted panels). **B) Left:** Both increasing groups exhibited better late-life verbal memory than the consistently low group (* p < .05 and ** p < .01), although the low-increasing group remained inferior to the consistently high group (# p < .05). **Middle:** Both increasing groups had lower regional white matter volume than the consistently high group (# p < .05, ## p < .01), possibly reflecting missed opportunities for optimisation during midlife. **Right:** The low-increasing group showed a cortical free water profile more similar to the consistently low group than to the consistently high group (# p <.05), whereas the average-increasing group exhibited lower free water compared with the consistently low group (* p < .05) while not statistically differentiable from the consistently high group. Y axes in panel B represent estimated marginal means of respective measures adjusted for key covariates using the R emmeans package. Error bars represent ±1 standard error of the group mean. Abbreviations: Mid = Midlife, Early = Early elderhood, Late = Late life.

Focusing on the effect of diet improvement (details in **Supplementary Results – DASH trajectory analyses**), we compared the low-increasing and average-increasing groups with the least favourable (consistently low) and most favourable (consistently high) trajectories across late life brain and cognitive measures identified above. We found that participants in the low-increasing group demonstrated better verbal memory than the consistently low group (t = 2.13, p = .034, p_fdr = .042), although they remained inferior to the consistently high group in verbal memory (t = −2.28, p = .023, p_fdr = .038), white matter volume (t = −2.86, p = .004, p_fdr = .007) and cortical free water (t = 2.42, p = .016, p_fdr = .02). In contrast, participants in the average-increasing group exhibited better verbal memory (t = 2.92, p = .004, p_fdr = .009), a trend towards greater white matter volume (t = 1.88, p = .061, p_fdr = .10), and lower cortical free water (t = −2.32, p = .02, p_fdr = .051) compared with the consistently low group. Notably, although this group had lower white matter volume than the consistently high group (t = −2.16, p = .031, p_fdr = .039), they were comparable in verbal memory and cortical free water (**Fig. 6B**). Collectively, these findings demonstrate meaningful benefits of post-midlife dietary improvement for late-life cognitive and brain health outcomes.

### Main results were robust across all validation analyses

The identified associations of life-stage diet with verbal memory and brain measures (i.e., white matter volume, deep basal ganglia WMH volume, and free water) were robust to progressive covariate adjustment from time-invariant demographic factors in Model 1, to the inclusion of time-specific health and lifestyle covariates in Model 2, and further adjustment for late-life health and lifestyle factors in Model 3 (**Methods – Statistical analyses – Multiple regression models)**, with the exception of grey matter morphometry and anterior periventricular WMH volume (**Supplementary Results – Cognitive performance, White matter morphometry, Grey matter morphometry, Regional white matter hyperintensities, and Brain free water**).

We further repeated the same analyses in a subset of healthier participants (N = 788) without suspect cognitive impairment and depressive symptoms (see **Methods**) and all key findings remained (**Supplementary Results – Validation analyses**).

## DISCUSSION

In this life-course study of Asian older adults with repeated dietary assessments alongside comprehensive neuropsychological testing and multimodal MRI, we address key limitations of prior studies to evaluate the enduring influence of diet across the life course on late-life cognitive health ^8,11,12^ in a domain-specific manner, to focus on the impact of dietary change over time, and to identify candidate neurobiological substrates linking diet to cognitive resilience ^3,18^. Higher past adherence to the DASH dietary pattern was associated with better verbal memory, greater white matter volume, lower cortical free water, and lower regional WMH in late life. Importantly, these brain phenotypes partially mediated the cognitive benefits associated with diet. Trajectory analyses further indicated that diet improvements after midlife were still associated with more favourable late-life cognitive and brain outcomes. Collectively, these findings advance our understanding of how life-course dietary quality modulate cognitive and brain health in late life.

Exemplifying the impact of DASH on cognitive health, a recent large-scale study (N = 159,347) reported a robust association between midlife DASH adherence and both late-life cognitive performance and subjective cognitive decline ^11^. Our findings further suggest domain-specific effects, with verbal memory emerging as the domain most sensitive to long-term dietary quality. This selectivity is not only consistent with reported vulnerability of verbal memory to cardiovascular and metabolic health ^8,38^, both closely associated with dietary patterns ^39^, but also its status as one of the earliest domains to decline prior to overt cognitive impairment ^40^, outperforming hippocampal atrophy and visuospatial measures in predicting progression to mild cognitive impairment ^41^. Critically, midlife diet quality remained independently associated with verbal memory more than two decades later, suggesting that sustained adherence to a healthy dietary pattern from midlife may modulate the cardiometabolic risk factors that preferentially threaten this domain, potentially preserving a function with clinical relevance.

A healthy diet may dynamically modulate neurobiological plasticity across adulthood, mitigating age-related degenerative processes and supporting cognition ^7,42^. Our dietary assessment by life stages allows us to extend prior cross-sectional and cohort studies with single dietary assessment ^13^ to suggest that associations with late-life white matter volume are most pronounced when diet is assessed in midlife, a period when accelerated white matter loss could prelude mild cognitive impairment ^43^. Healthy dietary patterns may slow this trajectory by supporting axonal integrity and vascular health, thereby contributing to structural brain reserve in later life ^44^. Importantly, the observed partial mediation suggests that midlife diet contributes to the preservation of white matter, which in turn supports late-life verbal memory. Consistent with this, these white matter clusters encompass brain networks implicated in language and verbal processing ^34^, and our previous work linked accelerated frontal and parietal white matter atrophy to faster verbal memory decline in ageing ^33^.

In grey matter, higher DASH adherence in early and late elderhood was associated with more favourable macrostructural measures in the precuneus, somatomotor cortex, and inferior prefrontal regions – areas that overlap with a putative lexico-semantic network supporting the adaptive reorganization of cognitive processing in older adults ^45^. As these regions typically mature by midlife before undergoing gradual decline ^45^, it is plausible that healthy dietary adherence after midlife may help attenuate this trajectory. However, the variability of grey matter findings across analytical models warrants cautious interpretation.

In contrast to macrostructural white matter measures, cortical grey matter free water (FW), a measure that reflects extracellular fluid alterations linked to neurovascular inflammation ^24,35,46^ and early ageing and cognitive impairment ^47^ but is underexplored in dietary research, was inversely associated with DASH adherence in early elderhood. This novel relationship was localised to left frontal and right occipital regions, with a more widespread cortical distribution observed in late life. The observed partial mediation suggests a complementary neurobiological correlate whereby healthier diet may preserve late-life verbal memory by attenuating neurovascular inflammatory burden ^3,18^. This possibility is supported by emerging evidence linking neuroinflammatory and neuroimmune processes to verbal memory impairment ^48^, and plausibly involving the gut microbiome ^49^. For the free water topography, our recent work has linked free water alterations in both anterior and posterior regions to poorer cardiovascular health and memory performance ^35,36^. Consequently, healthier dietary patterns after midlife may attenuate neurovascular inflammatory burden in these regions ^50^, potentially offering resilience to neural functioning.

While structural reserve may be consolidated in midlife, the mitigation of neurovascular inflammation may become increasingly relevant later in life, when degenerative processes accelerate ^7^ and the consequences of poor dietary habits are more acutely expressed ^50^. Since we only collected MRI data in late life, future studies with longitudinal neuroimaging data across the life course are needed to further verify these age-dependent associations. We also note that both observed mediation effects were modest in magnitude, consistent with prior neuroimaging ^27^ and lifestyle studies ^51^. Examining additional diet-sensitive brain measures ^52,53^, and refining assessments beyond broad dietary patterns ^54^ and cognitive constructs ^55^, will be important for clarifying the relationship between diet and healthspan.

Although white matter free water is sensitive to ageing and cardiovascular risk ^35,36^, no significant associations with diet were observed, warranting further investigation. In contrast, white matter hyperintensities (WMHs), a marker of cerebral small vessel disease and closely related to free water ^56^, were negatively associated with DASH adherence in early elderhood. Prior studies have linked regional WMHs around the deep basal ganglia and anterior periventricular regions to vascular risk factors ^26^ and vascular dementia ^57^, suggesting that both aetiology (e.g., vascular health) and spatial distribution should be considered when exploring diet-WMH associations. The vascular involvement may be particularly relevant in Asian populations, where cardiometabolic risk profiles differ from those of Western populations ^58,59^. Targeted dietary strategies such as DASH may play a meaningful role in reducing cardiometabolic and dementia risk in this context ^60,61^. Interestingly, in contrast to some non-dietary studies ^62^, regional WMHs were not significantly associated with cognition in the present analysiss, highlighting the need for further investigation of how diet and lifestyle shape cerebrovascular and cognitive outcomes ^63,64^.

Finally, our trajectory analyses identified subgroups of older adults who improved their diet quality from low or average levels after midlife and subsequently demonstrated better late-life verbal memory than those who maintained persistently poor dietary patterns, extending our previous finding on cognitive impairment to a cognitive domain that is highly clinically relevant ^5^. Notably, individuals in the average-increasing group achieved verbal memory performance comparable to those with consistently high diet quality. Improvements in late-life brain measures followed the life-stage patterns (i.e., missed opportunities on white matter volume consolidation in midlife) and were more modest, although participants in the average-increasing group exhibited lower cortical free water than those in the consistently low group and was comparable to those in the consistently high group, suggesting some neurobiological benefit even from late-initiated dietary improvement ^65^. We consider these findings to have practical relevance since most people maintain diets of average rather than exemplary quality. Together, our results reinforce the importance of sustaining high diet quality throughout adulthood ^66^, while also highlighting a potential window for effective dietary intervention after midlife ^67^. The magnitude and nature of benefit likely depend on which neurobiological substrates remain responsive to dietary and nutritional influences at different life stages. Realising the potential of this window will require addressing critical issues such as how dietary protective effects can be fully harnessed amid age-related changes in appetite, energy needs, and eating behaviour ^6^, and how intervention trials or public health programs can balance individual effectiveness with scalability across diverse ageing populations ^18^.

We acknowledge several limitations. First, survivorship bias, recall error, and social desirability bias inherent to self-reported dietary intake may be present; however, these are more likely to attenuate observed associations ^12^. Second, although different dietary questionnaires were administered across study visits, DASH scores were derived using harmonised food items ^12^, and associations between life-stage DASH adherence and late-life verbal memory remained consistent. Third, while we adjusted for key lifestyle factors, including physical activity, future studies should examine potential interactions between diet and other behaviours ^68,69^. Finally, in addition to neuroimaging, longitudinal cognitive data are required to fully characterize the relationship between intra-individual dietary and neurocognitive trajectories.

In conclusion, we investigated diet, a modifiable and scalable lifestyle factor, in relation to brain and cognitive health from a life-course perspective, leveraging the detailed historical diet, neuroimaging, and cognitive assessments from a large group of Asian older adults in the SG70 cohort. We show that higher diet quality across adulthood is associated with preserved late-life verbal memory, potentially mediated by greater structural brain reserve and reduced neurovascular inflammatory burden. Notably, improvements in diet after midlife were also associated with more favourable late-life cognitive and brain health outcomes. These findings advance our understanding of neurocognitive ageing and inform both clinical practice and public health policy with respect to healthy diet promotion.

## METHODS

### Participants

The ongoing population-based Singapore Chinese Health Study (SCHS) was first established to investigate the determinants of common diseases in Asian populations ^29^. A total of 63,257 Singaporean Chinese adults aged 44-74 years, free of dementia at baseline, were recruited between 1993 and 1998 (midlife). Surviving and consenting participants were subsequently followed up in 1999-2004 (follow-up 1), 2006-2010 (follow-up 2), and 2014-2016 (follow-up 3). Between 2021 and 2025 (late life; follow-up 6), 1,165 participants returned as part of the ‘SG70: Towards Healthy Longevity’ cohort ^30^ (aged 68-82 years), which was dedicated to the study of ageing outcomes among participants in their seventies, giving rise to the subcohort name SG70. Among a subset of SG70 participants who underwent multimodal magnetic resonance imaging (MRI) and domain-specific neuropsychological assessments, 892 individuals (489 women) with largely complete MRI, dietary, and neuropsychological assessment data (small proportion of missing demographic and assessment information) were included in the present analysis (**Table 1**). All procedures in the SCHS and SG70 studies were approved by the Institutional Review Board of the National University of Singapore and the Domain Specific Review Board (DSRB) of the National Healthcare Group (NHG Health). All participants provided written informed consent.

### Diet quality assessment

Diet quality at each time point was assessed using adherence to a modified Dietary Approaches to Stop Hypertension (DASH) pattern. Briefly, at baseline, dietary intake in the past one year was assessed using an interviewer-administered, 165-item, semi-quantitative food frequency questionnaire (FFQ), in which participants indicated their frequency of consumption for each food item ^12^. Energy intake per day (kcal) was also estimated. At follow-up 3 and SG70 study, dietary intake in the past one year was assessed using a modified dietary screener on the consumption of 21 food and beverage items, which provided sufficient information for DASH derivation. Notably, total dairy intake was taken as a surrogate for the calculation of the low-fat dairy component, and the sodium component was excluded due to restrictions of the dietary instruments.

### Neuropsychological assessment

A detailed neuropsychological assessment battery was developed based on our previous work ^32,33,70^ and administered by trained researchers. A total of nine tests covered seven major cognitive domains. Specifically, processing speed was assessed using the Symbol Digit Modality Test and the Colour Trail Making Test A. Attention was assessed using the Forward and backward Digit Span Task. Executive function was assessed using the Design Fluency Test in the Delis-Kaplan Executive Function System and the Colour Trail Making Test B. Visuospatial ability was assessed using the Block Design Test. Visual memory was assessed using the Paired Associative Learning task in CANTAB and Brief Visuospatial Memory Test–

Revised. Verbal memory was assessed using the Rey Auditory Verbal Learning Test (RAVLT). Finally, language ability was assessed using a modified Boston Naming Task (BNT) for Asian cohorts. For the current study, individual test scores were standardised as T scores (mean = 50, standard deviation = 10) and inverted if needed, such that a higher score always indicated better performance. Domain composite scores were then computed by averaging the T scores from relevant tests. This procedure yielded seven cognitive domain composite scores for each participant.

To further stratify participants according to their mental health and cognitive status, we administered the 15-item Geriatric Depression Scale (GDS) and the Quick Dementia Rating System (QDRS), the scores from which were converted to the Clinical Dementia Rating (CDR) based on the criteria from the original article. See **Supplementary Methods** for detailed references to these tests.

### MRI acquisition and preprocessing

MRI data were collected in a 3T Siemens Prisma-fit scanner at the Centre for Translational MR Research in the National University of Singapore using a 32-channel head coil. Anatomical images were collected with a T1-weighted Magnetisation Prepared Rapid Gradient Recalled Echo sequence (MPRAGE; 176 sagittal slices, resolution 1 mm isotropic, repetition time = 1950 ms, echo time = 2.98 ms). T2-FLAIR images were obtained with a 3D fast spin-echo sequence (SPACE; 192 sagittal slices, resolution 1 mm isotropic, repetition time = 7000 ms, echo time = 393 ms). Diffusion-weighted images were collected with a modified multi-shell, spin-echo EPI protocol ^71^ (TR/TE = 3200/79.4 ms, FOV = 216 mm, 81 contiguous slices, voxel size = 1.8 mm^3^ isotropic, multi-band factor = 3), using interleaved shells at b = 200, 500, 1000, and 2000 s/mm² (6, 10, 50, and 50 directions, respectively) and 8 b0 images. Reversed phase-encoding b0 blocks (Anterior-Posterior) were additionally acquired for susceptibility-distortion correction (6 volumes before and 6 volumes after the main Posterior-Anterior run). Of note, one b = 3000s/mm^2^ volume was included in both runs to ensure scanning parameters consistency with the original protocol; they were excluded from further preprocessing.

Anatomical T1 data were pre-processed using FreeSurfer V5.3 (https://surfer.nmr.mgh.harvard.edu) and Computational Anatomy Toolbox (CAT12) (https://neuro-jena.github.io/cat/) separately to obtain both vertex-and voxel-level measurements. For FreeSurfer, steps included motion correction, removal of the non-brain mass, automated Talairach transformation, intensity correction, volumetric segmentation, cortical surface parcellation, and reconstruction. For CAT12, steps included bias-field correction, denoising, skull-stripping, tissue classification, spatial normalisation to the MNI template, and generation of modulated tissue probability maps that preserve regional volume information. FLAIR data were minimally preprocessed using the FSL Brain Extraction Tool followed by visual QC to extract all brain tissues. Finally, diffusion MRI data were pre-processed using FSL V6.0 (http://www.fmrib.ox.ac.uk/fsl) with steps including eddy current correction, diffusion gradient rotation, and geometric distortions correction. All data were checked for data quality, co-registration quality, and excessive motion.

### MRI measures derivation

#### Brain morphometry

Vertex-wise morphometric maps of cortical volume, thickness, and surface area were generated at the final step of the FreeSurfer preprocessing. Voxel-wise grey matter volume (GMV) and white matter volume (WMV) were generated at the final step of the CAT12 preprocessing. Spatial smoothing with Gaussian kernels of 15 mm and 8 mm full width at half maximum (FWHM) were applied to vertex-wise and voxel-wise data, respectively, for each measure and participant.

#### Regional white matter hyperintensities (WMH)

WMH was segmented from FLAIR images using WMH-DualTasker, a weakly supervised deep-learning based algorithm we developed recently ^72^, followed by manual checking. To obtain regional WMH patterns, the Bullseye segmentation ^26^ was applied to create 36 WM parcels (4 equidistant zones between the ventricular surface and the white matter surface x 4 lobes x 2 hemispheres + 4 equidistant basal ganglia zones). Parcellated WMHs were then subject to the exploratory graph analysis (EGA), a graph-based factor analytic method. This procedure yielded seven WMH spatial patterns that were labelled according to their spatial distributions: anterior periventricular, frontoparietal, white matter around deep basal ganglia, left and right (occipital) horns, temporal, and parietal ^26^. The regression method was used to extract a factor score for each spatial pattern and participant.

#### Brain free water (FW)

FW maps were derived from preprocessed diffusion MRI data using FISO maps from the NODDI model ^46^, together with a free water imaging approach adapted from our previous work ^35,36^. For cortical grey matter, FISO maps were projected onto the cortical surface to generate vertex-wise maps with partial volume correction. Subcortical grey matter FISO maps were retained in volumetric space. In both cortical and subcortical regions, analyses were restricted to vertices or voxels with a grey matter tissue probability of at least 30% ^35,36^. For white matter, volumetric FISO maps were further processed using tract-based spatial statistics (TBSS) to generate skeletonised maps aligned to a group-level white matter tract space for each participant. The skeleton template was created based on participants’ FA maps (b = 1000) with a threshold of 0.2.

### Statistical analyses

#### Multiple regression models

To examine the associations between life-stage diet quality and late-life cognitive and brain measures, we implemented multiple regression models following our previous approach ^12^. DASH adherence at each time point was modelled separately as the exposure of interest, with demographic, self-reported health conditions, and lifestyle covariates progressively included across three models. Model 1 served as the base (“discovery”) model and included core covariates: daily energy intake (measured at midlife), age at follow-up 6, sex, handedness, dialect group, and education. Model 2 extended by additionally adjusting for health and lifestyle variables measured at the same time point as the DASH exposure. For midlife DASH, these included amount of physical activity at work and sport, self-report medical conditions (cancer, stroke, type 2 diabetes, hypertension, and any heart conditions), alcohol consumption, smoking, body mass index (BMI) categories, and average sleep duration per day. For early elderhood DASH, corresponding measures included amount of physical activity, new cases of medical conditions, alcohol consumption, smoking, BMI categories, and average sleep duration per day. For late-life DASH, corresponding measures included current medical conditions, current alcohol and smoking, BMI categories, and average sleep duration per day ^30^. Finally, Model 3 further extended Model 2 by incorporating health and lifestyle covariates measured at follow-up 6. For late-life DASH, Model 2 and 3 were therefore the same. For all brain-related analyses (except cortical thickness), intracranial volume was included as a covariate in all models.

#### Associations between diet history and cognition

To evaluate if life-stage diet quality was associated with late-life cognitive performance, we applied Model 1 with each of the seven cognitive domain composite scores as outcome separately, adjusting for multiple comparisons of the seven domains using False Discovery Rate (q = .05). Cognitive domains showing statistically significant effects with DASH adherence were further evaluated using Model 2 and Model 3. For cognitive domains significantly linked to midlife DASH adherence, to further examine if midlife diet quality has an enduring influence on late-life cognition, we performed an additional multiple regression analysis by including all DASH scores in Model 3.

#### Associations between diet history and brain morphometry

To examine associations between life-stage diet quality and late-life brain macrostructure, we performed mass-univariate multiple regression analyses on vertex and volumetric measures. Vertex-wise general linear models (GLMs) were applied to cortical thickness, surface area, and volume using the VertexWiseR package, while voxel-wise GLMs were applied to grey and white matter volume maps using the CAT12 GLM module. Cluster level inferences were used for both vertex- and voxel-wise analyses, with a cluster forming threshold of p = .01 and cluster-level significance of p = .05, familywise error (FWE) corrected. Statistically significant clusters were first identified using Model 1. Robustness was then evaluated in two ways. First, the vertex- and voxel-wise GLMs were repeated with Model 2 and Model 3 to assess spatial consistency of the effects. Second, mean values within clusters identified in Model 1 were aggregated and further evaluated with Model 2 and Model 3 using Bayesian multiple regression models with the brms package. Robustness was defined by consistent effect size estimates across models, with all corresponding 95% credible intervals excluding zero. These results were documented in **Supplementary Results – White matter morphometry** and **Grey matter morphometry**.

For anatomical interpretation, significant clusters from grey matter morphometry were mapped onto the Schaefer’s functional atlas ^37^, which delineated eight canonical resting-state networks (default mode, control, salience/ventral attention, dorsal attention, visual, somatomotor, limbic, or temporoparietal). Significant clusters from white matter morphometry were mapped onto population-based tractography atlas derived from healthy adults (HCP-YA1065) ^73^ and the WhiteRest Atlas that parcellated grey and white matter jointly into brain networks ^34^.

#### Associations between diet history and regional WMH

To examine associations between life-stage diet quality and late-life WMHs burden, the factor scores of the seven WMH spatial components were subject to multiple regression analyses at each time point using Model 1, adjusting for multiple comparisons of seven spatial components (FDR q = .05). Statistically significant components were further assessed with Model 2 and Model 3. These results were documented in **Supplementary Results – Regional WMH.**

#### Associations between diet history and brain free water

To examine associations between life-stage diet quality and late-life brain free water (FW), mass-univariate GLMs were applied to grey and white matter free water maps. Vertex-wise analyses of cortical grey matter FW were conducted using an in-house GLM framework based on FreeSurfer Monte-Carlo cluster-wise inference (mri_glmfit; 10,000 iterations; p<0.05 FWE corrected) ^35,36^. Voxel-wise analyses of subcortical grey matter FW were performed using the CAT12 GLM module as the morphometry analyses. Skeletonised white matter FW maps were analysed using FSL Randomise (threshold-free cluster enhancement, 5,000 permutations). As in the macrostructural analyses, statistically significant clusters were identified using Model 1, followed by robustness evaluation. These results were documented in **Supplementary Results – Brain free water.** For anatomical interpretation, significant grey matter clusters were mapped onto the Schaefer cortical atlas and the AAL subcortical atlas, whereas white matter clusters were mapped onto the HCP-YA1065 tractography atlas.

#### Association between brain measures and cognition

To evaluate the functional relevance of identified brain measures, multiple regression analyses were conducted relating cognitive performance to both DASH adherence and brain measures (cognitive score ∼ DASH + brain measure + covariates) across Models 1–3. Analyses were restricted to cognitive domains and brain measures with significant associations with diet quality. For vertex- and voxel-wise measures, mean values were extracted from significant clusters identified in Model 1.

#### Mediation analyses

To examine whether diet-sensitive brain measures may serve as neurobiological mediators, we conducted mediation analyses using path models implemented in the lavaan package. To focus on the effects of past diet, analyses were restricted to cognitive and brain measures that showed significant associations with DASH adherence at baseline or early elderhood. For vertex- and voxel-wise brain measures, mean values were extracted from significant clusters identified in Model 1. All variables included in the path models were residualised with respect to demographic, health and lifestyle covariates in Model 3. Statistical significance of the indirect effects (i.e., diet brain cognition) was determined using the 95% confidence intervals derived from 5,000 bootstrapping. Percentage of mediation was computed as the proportion of the indirect effect over the total effect.

#### DASH trajectory derivation and analyses

To evaluate whether improvements in diet after midlife confer benefits to late-life brain and cognitive health ^12^, we performed a latent class growth analysis (LCGA) on DASH adherence across all time points using the lcmm package, restricting to linear trajectory and within-group invariance ^74^. Mean age of each time point was used as the metric of time and normalised as [0, .72, 1] ^75^. Models with one to seven classes were compared based on AIC, BIC, and sample size adjusted BIC (saBIC) alongside class interpretability (**Supplementary Results – DASH trajectory analyses**). To ensure global convergence, each model was fitted 200 times with different initialisation. Trajectory groups were defined based on estimated intercepts (low, average, high midlife adherence) and slopes (consistent, increasing, decreasing), and ordered from least to most favourable. Participants were assigned to groups based on class posterior probabilities.

Subsequently, multiple regression analyses were conducted to examine associations between trajectory group membership and brain and cognitive measures implicated in the mediation analyses. Group effects were first ascertained using omnibus F-tests (**Supplementary Results – DASH trajectory analyses**). Selective pairwise contrasts comparing improving groups with the least favourable (consistently low) and most favourable (consistently high) groups were then examined using Wald tests. All analyses adjusted for key covariates measured at midlife and late life. For a complete understanding of the diet trajectory influences, we provided the summary of the remaining non-improving groups and statistical significance adjusting for all pairwise comparisons with each anchoring group in **Supplementary Results – DASH trajectory analyses.**

#### Validation analyses

To show that our diet effects were not due to the inclusion of participants who might have atypical cognitive or mental health deficits, associations between DASH adherence, cognition, and brain measures were re-examined in a subgroup of ‘healthier’ participants by excluding those (N = 104) who failed more than 3 cognitive domains (1.5 SD below Singaporean norms)^76^ (N = 12), showed depressive symptoms (GDS >= 5; N = 63), or had a non-zero CDR score based on QDRS (N = 50). These results were documented in **Supplementary Results – Validation analyses.**

## Supporting information

Supplementary Methods and Results

## Data availability

The datasets generated and/or analysed during the current study are not publicly available due to data privacy laws, but they are available from the corresponding authors on reasonable request.

## Code availability

WMH-DualTasker is publicly available in https://github.com/yilei-wu/WMH-DualTasker. Codes for WMH Bullseye parcellation are publicly available in https://github.com/gsanroma/bullseye_pipeline. Exploratory Graph Analysis was conducted using the EGA package (https://r-ega.net/). Vertex-wise mass univariate analyses were conducted using the VertexWiseR package (https://github.com/CogBrainHealthLab/VertexWiseR). Mediation analyses were conducted using the lavaan package (https://lavaan.ugent.be/). LCGA was conducted using the lcmm package (https://cecileproust-lima.github.io/lcmm/). Marginal means were evaluated using the emmeans package (https://cran.r-project.org/web/packages/emmeans/index.html). R codes used to conduct main analyses are available in https://github.com/hzlab/2026_Ng_SG70_diet

## Acknowledgements

The SCHS was supported by grants from the Singapore National Medical Research Council (NMRC/CSA/0055/2013, Singapore Strategic Cohorts Consortium Award P2022-02) and the U.S. National Institutes of Health (grant numbers R01 CA144034, and UM1 CA182867). The SG70 project was supported by Singapore National Medical Research Council [CSA-SI (MOH-000434)], Yong Loo Lin School of Medicine, National University of Singapore, as well as grants from the Singapore National Medical Research Council (HLCA23Feb-0004, OFIRG24Jul-0049, and HPJGC25-0002).

We thank Ms. Siew-Hong Low of the National University of Singapore (NUS) for overseeing the fieldwork of the Singapore Chinese Health Study and SG70; Dr. Ruth Li Fang Leong of NUS and Mr. Eddie Jun Yi Chong of the National University Hospital (NUHS) for consultation on the development of the neuropsychological assessment battery; Dr. Siwei Liu of NUS for consultation on diffusion MRI acquisition, and Ms. Vellica Filbert, Ms. Claudia Mei Sha Ho, Ms. Sherlyn Pei Xin Tang, and Ms. Seetha Harinakshi Sattanathan of NUS for assisting in the WMH segmentation.

## Author contributions

E.K.K.N., J.S.L., W.-P.K., and H.J.Z conceived, conceptualized and designed the study. E.K.K.N., J.S.L., and F.J. analysed the data. Y.L.B.G., P.S.Q.A., W.L.L., and J.S.Y.T collected, preprocessed, and managed the data. E.K.K.N. wrote the first draft of the paper. E.K.K.N., S.F., W.-P.K., and H.J.Z. interpreted the findings and wrote the revised versions of the paper.

