## Supplementary Methods and Results for "Improving DASH adherence after midlife promotes better late-life cognition and brain health: the SG70 Study"

A. **Supplementary Methods**

A total of nine tests covered seven major cognitive domains. Specifically, processing speed was assessed using the Symbol Digit Modality Test ^1^ and the Colour Trail Making Test A ^2^. Attention was assessed using the Forward and backward Digit Span Task ^3^. Executive function was assessed using the Design Fluency Test in the Delis-Kaplan Executive Function System ^4^ and the Colour Trail Making Test B ^5^. Visuospatial ability was assessed using the Block Design Test ^3^. Visual memory was assessed using the Paired Associative Learning task in CANTAB ^6^ and Brief Visuospatial Memory Test–Revised ^7^. Verbal memory was assessed using the Rey Auditory Verbal Learning Test (RAVLT) ^8^. Finally, language ability was assessed using a modified Boston Naming Task (BNT) for Asian cohorts ^9^. An example of the T-score conversion can be found in ^10^. We also administered the 15-item Geriatric Depression Scale (GDS) ^11^ and the Quick Dementia Rating System (QDRS)^12^, the scores from which were converted to the Clinical Dementia Rating (CDR ^13^) based on the criteria from the original article.

WMH-DualTasker ^14^ is publicly available in <https://github.com/yilei-wu/WMH-DualTasker>. WMH bullseye representation was first proposed by Sudre and colleagues ^15^ and implementation codes are publicly available in <https://github.com/gsanroma/bullseye_pipeline>. Exploratory Graph Analysis was conducted using the EGA package ^16^ (https://r-ega.net/). Vertex-wise mass univariate analyses were conducted using the VertexWiseR package ^17^ (https://github.com/CogBrainHealthLab/VertexWiseR). Bayesian multiple regression modelling was performed using the brms package ^18^ (https://github.com/paul-buerkner/brms). Mediation analyses were conducted using the lavaan package ^19^ (<https://lavaan.ugent.be/>). LCGA was conducted using the lcmm package ^20^ (<https://cecileproust-lima.github.io/lcmm/>). Marginal means were evaluated using the emmeans package ^21^.

B. **Supplementary Results – Cognitive performance**

DASH adherence in midlife, early elderhood, and late life was most consistently associated with late-life verbal memory, after adjustment for multiple comparisons across seven cognitive domains in each life stage (**Supplementary Table 1**). These associations remained following additional covariate adjustments (**Supplementary Table 2**). In addition, midlife DASH adherence retained an independent association with late-life verbal memory when all life-stage DASH scores were modelled jointly (**Supplementary Table 3**).

*Supplementary Table 1. Associations between higher life-stage DASH adherence and better late-life verbal memory from Model 1, controlling for basic demographic covariates*

| DASH | Cognitive domain | *b* | *SE* | *t* | *p* | p_fdr | 95% CI |
| --- | --- | --- | --- | --- | --- | --- | --- |
| Midlife | **Verbal memory** | **0.19** | **0.06** | **3.18** | **.002**** | **.011*** | **[0.07, 0.31]** |
|  | Visual memory | -0.04 | 0.06 | -0.68 | .495 | 1.00 | [-0.15, 0.07] |
|  | Visuospatial ability | -0.02 | 0.07 | -0.33 | .740 | 1.00 | [-0.15, 0.11] |
|  | Executive functioning | -0.05 | 0.06 | -0.86 | .388 | 1.00 | [-0.16, 0.06] |
|  | Attention | -0.03 | 0.06 | -0.51 | .608 | 1.00 | [-0.15, 0.09] |
|  | Processing speed | 0.02 | 0.05 | 0.39 | .696 | 1.00 | [-0.08, 0.12] |
|  | Language | -0.01 | 0.06 | -0.16 | .874 | 1.00 | [-0.13, 0.11] |
| Early elderhood | **Verbal memory** | **0.20** | **0.06** | **3.37** | **.001***** | **.006**** | **[0.08, 0.32]** |
|  | Visual memory | 0.09 | 0.06 | 1.64 | .101 | .505 | [-0.02, 0.20] |
|  | Visuospatial ability | 0.08 | 0.07 | 1.17 | .241 | .962 | [-0.05, 0.21] |
|  | Executive functioning | -0.02 | 0.06 | -0.34 | .732 | 1.00 | [-0.13, 0.09] |
|  | Attention | -0.05 | 0.06 | -0.77 | .444 | 1.00 | [-0.16, 0.07] |
|  | Processing speed | -0.04 | 0.05 | -0.86 | .389 | 1.00 | [-0.15, 0.06] |
|  | Language | 0.13 | 0.06 | 2.08 | .037* | .224 | [0.01, 0.25] |
| Late life | **Verbal memory** | **0.24** | **0.06** | **4.21** | **< .001***** | **< .001***** | **[0.13, 0.36]** |
|  | Visual memory | 0.10 | 0.05 | 1.93 | .054 | .283 | [-0.00, 0.20] |
|  | Visuospatial ability | 0.07 | 0.06 | 1.22 | .223 | .777 | [-0.05, 0.19] |
|  | Executive functioning | 0.07 | 0.05 | 1.30 | .194 | .777 | [-0.04, 0.17] |
|  | Attention | -0.04 | 0.05 | -0.75 | .454 | .908 | [-0.15, 0.07] |
|  | Processing speed | 0.03 | 0.05 | 0.64 | .522 | .908 | [-0.06, 0.13] |
|  | Language | 0.11 | 0.06 | 1.99 | .047* | .283 | [0.00, 0.23] |

*Note.* Model 1 controlled for total food energy measured in baseline, age at follow-up 6, handedness, sex, dialect group, and levels of education. b = beta coefficient, SE = standard error, CI = 95% confidence intervals, p_fdr = multiple comparison correction across seven domains. * p < .05, ** p < .01, *** p < .001

| *Supplementary Table 2. Association between DASH adherence and verbal memory at late life adjusting for additional time-specific (Model 2) and time-specific and late-life (Model 3) covariates* | | | | | | | |
| --- | --- | --- | --- | --- | --- | --- | --- |
| DASH | Model | Cognitive domain | *b* | *SE* | *t* | *p* | 95% CI |
| Midlife | 2 | Verbal memory | 0.23 | 0.07 | 3.57 | < .001*** | [0.10, 0.36] |
| Early elderhood | 2 | Verbal memory | 0.21 | 0.06 | 3.34 | .001*** | [0.09, 0.33] |
| Late life | 2 | Verbal memory | 0.24 | 0.06 | 3.77 | < .001*** | [0.11, 0.36] |
| Midlife | 3 | Verbal memory | 0.21 | 0.06 | 3.32 | .001*** | [0.09, 0.34] |
| Early elderhood | 3 | Verbal memory | 0.20 | 0.06 | 3.22 | .001** | [0.08, 0.33] |
| *Note:* Model 1 controlled for total food energy measured in baseline, age at follow-up 6, handedness, sex, dialect group, levels of education, and intracranial volume; Model 2 controlled for Model 1 and key covariates measured at the same time point as DASH. Model 3 controlled for Model 2 and key covariates measured at follow-up 6. b = beta coefficient, SE = standard error, CI = 95% confidence intervals, * p < .05, ** p < .01, *** p < .001 | | | | | | | |

| *Supplementary Table 3. Association between diet and verbal memory considering all DASH scores jointly, controlling for key covariates in midlife and late life (Model 3)* | | | | | | |
| --- | --- | --- | --- | --- | --- | --- |
| DASH | Cognitive domain | *b* | *SE* | *t* | *p* | 95% CI |
| Midlife | Verbal memory | 0.16 | 0.07 | 2.32 | .021* | [0.02, 0.30] |
| Early elderhood | Verbal memory | 0.07 | 0.08 | 0.94 | .345 | [-0.08, 0.22] |
| Late life | Verbal memory | 0.15 | 0.08 | 1.98 | .048* | [0.00, 0.30] |
| *Note.* b = beta coefficient, SE = standard error, CI = 95% confidence intervals. * p < .05, ** p < .01, *** p < .001 | | | | | | |

C. **Supplementary Results – White matter morphometry**

Midlife DASH adherence was associated with greater late-life white matter volume in inferior prefrontal, parietal, and cingulate areas (**Supplementary Table 4**). Clusters remained consistent across the three regression models with progressive covariate adjustment. Bayesian analyses of the supra-threshold clusters identified in Model 1 indicated consistent effect sizes of the diet-WMV association following additional covariate adjustment (**Supplementary Table 5**). Significant WMV clusters encompassed multiple white matter fibre tracts, many of which are association fibres and involved in language and verbal processes (**Supplementary Fig. 1**).

| *Supplementary Table 4. Cluster peak statistics of voxel-wise analysis between midlife DASH adherence and late life white matter volume (WMV) in the MNI space* | | | | | | |
| --- | --- | --- | --- | --- | --- | --- |
| DASH | Cluster | X | Y | *Z* | Size (voxels) | Peak Z value |
| Midlife | 1 | 38.00 | 45.00 | 3.00 | 2268 | 4.39 |
|  | 2 | 38.00 | -34.00 | 20.00 | 5779 | 4.31 |
|  | 3 | -51.00 | -50.00 | -3.00 | 3761 | 4.19 |

*Supplementary Table 5. Bayesian summary of midlife DASH association with white matter volume (WMV)*

| DASH | Model |  |  | Mean estimate | 95% Credible Interval |
| --- | --- | --- | --- | --- | --- |
| Midlife | 1 |  |  | .00109 | [.00060, .00157] |
|  | 2 |  |  | .00121 | [.00069, .00173] |
|  | 3 |  |  | .00111 | [.00058, .00164] |
| *Note:* Model 1 controlled for total food energy measured in baseline, age at follow-up 6, handedness, sex, dialect group, levels of education, and intracranial volume; Model 2 controlled for Model 1 and key covariates measured at the same time point as DASH. Model 3 controlled for Model 2 and key covariates measured at follow-up 6. | | | | | |

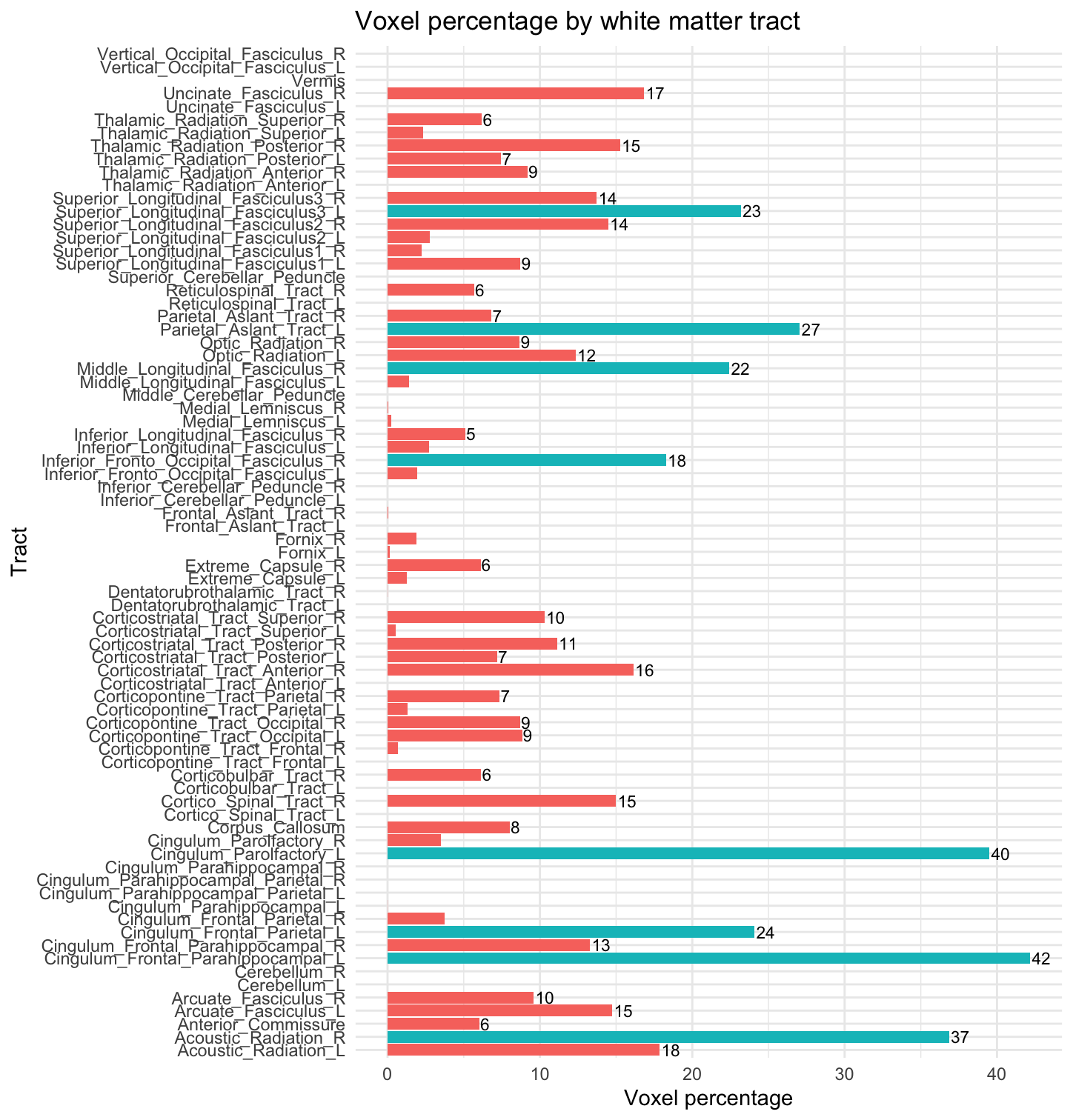

**Supplementary Figure 1.** Percentage overlap between major white matter fibre tracts ^22^ and statistically significant white matter volume clusters associated with higher DASH adherence in midlife (baseline). The top ten tracts are highlighted in turquoise for illustrative purpose.

D. **Supplementary Results – Grey matter morphometry**

DASH adherence in early elderhood and late life was associated with multiple macrostructural morphometry, including the precuneus, somatomotor cortices, and bilateral inferior frontal cortices **(Supplementary Table 6; Supplementary Fig. 2).** The association with precuneus thickness remained consistent across the three regression models with progressive covariate adjustment. Further, Bayesian analyses of the supra-threshold clusters identified in Model 1s supported consistent effect sizes for the diet-morphometry associations following additional covariate adjustment (**Supplementary Table 7**). For volumetric results, voxel-based (CAT12) and vertex-based (FreeSurfer) analyses did not yield concordant observations ^23^, and these results should therefore be interpreted with caution.

| *Supplementary Table 6. Cluster peak statistics of vertex-wise analysis between grey matter morphometry and DASH adherence at early elderhood and late life in MNI space* | | | | | | |
| --- | --- | --- | --- | --- | --- | --- |
| DASH | Measure | X | Y | *Z* | Size (vertices) | Peak Z value |
| Early elderhood | Thickness | -9.6 | -60.3 | 52.6 | 121 | 3.11 |
| Early elderhood | Volume | 41.2 | -32.7 | 50.5 | 173 | 5.79 |
| Late life | Volume | 45.3 | 8.5 | 20.7 | 147 | 3.55 |
| Late life | Volume | -44.2 | 28.6 | 13.3 | 158 | 4.27 |

| *Supplementary Table 7. Bayesian summary of the DASH effect on grey matter morphometry* | | | | | | |
| --- | --- | --- | --- | --- | --- | --- |
| DASH | Measure | Model |  |  | Mean estimate | 95% Credible Interval |
| Early elderhood | Precuneus Thickness | 1 |  |  | .64 | [.31, .97] |
|  |  | 2 |  |  | .63 | [.28, .98] |
|  |  | 3 |  |  | .68 | [.33, 1.04] |
|  | Somatomotor  Volume | 1 |  |  | 12.19 | [6.28, 18.05] |
|  |  | 2 |  |  | 9.97 | [3.77, 16.14] |
|  |  | 3 |  |  | 9.93 | [3.57, 16.12] |
| Late life | Bilateral inferior frontal Volume | 1 |  |  | 34.01 | [20.35, 47.45] |
|  |  | 2 (&3) |  |  | 34.45 | [19.52, 49.02] |
| *Note:* Model 1 controlled for total food energy measured in baseline, age at follow-up 6, handedness, sex, dialect group, levels of education, and intracranial volume; Model 2 controlled for Model 1 and key covariates measured at the same time point as DASH. Model 3 controlled for Model 2 and key covariates measured at follow-up 6. | | | | | | |

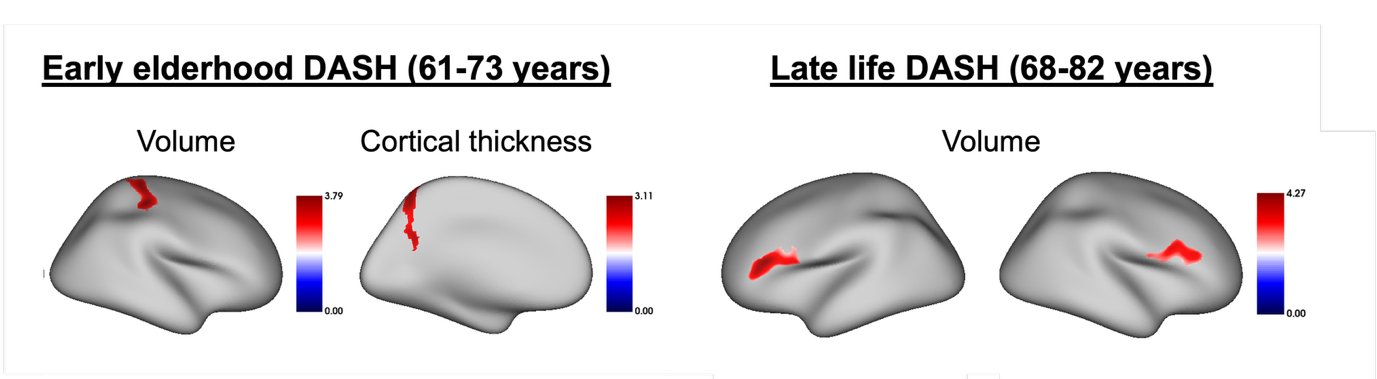

**Supplementary Figure 2.** Higher DASH adherence in early elderhood was associated with greater cortical thickness in the left precuneus and increased volume in somatomotor regions, whereas higher DASH adherence in late life was associated with larger bilateral inferior frontal volumes. Colour map represents t values.

E. **Supplementary Results – Brain free water**

Higher DASH adherence in early elderhood was associated with lower cortical grey matter free water in left frontal and right occipital regions, spanning attentional, default mode, control, and visual functional brain networks (**Supplementary Fig. 3**). Similar, but more spatially extensive, associations were observed cross-sectionally with late-life DASH adherence (**Supplementary Fig. 3 and 4, Supplementary Table 8**). All clusters remained consistent across the three regression models with progressive covariate adjustment. Bayesian analyses of the supra-threshold clusters identified in Model 1 indicated consistent effect sizes for the associations between DASH adherence and cortical grey matter free water following additional covariate adjustment (**Supplementary Table 8**).

*Supplementary Table 8. Bayesian summary of the DASH effect on cortical grey matter free water*

| DASH |  | Region | Model | Mean estimate | 95% Credible Interval |
| --- | --- | --- | --- | --- | --- |
| Early elderhood |  | cortex | 1 | -0.93 | [-1.34 -0.52] |
| Early elderhood |  | cortex | 2 | -0.85 | [-1.27, -0.43] |
| Early elderhood |  | cortex | 3 | -0.80 | [-1.22, -0.36] |
| Late life |  | cortex | 1 | -1.03 | [-1.42, -0.64] |
| Late life |  | cortex | 2 (&3) | -0.91 | [-1.36, -0.46] |

*Note:* Model 1 controlled for total food energy measured in baseline, age at follow-up 6, handedness, sex, dialect group, levels of education, and intracranial volume; Model 2 controlled for Model 1 and key covariates measured at the same time point as DASH. Model 3 controlled for Model 2 and key covariates measured at follow-up 6.

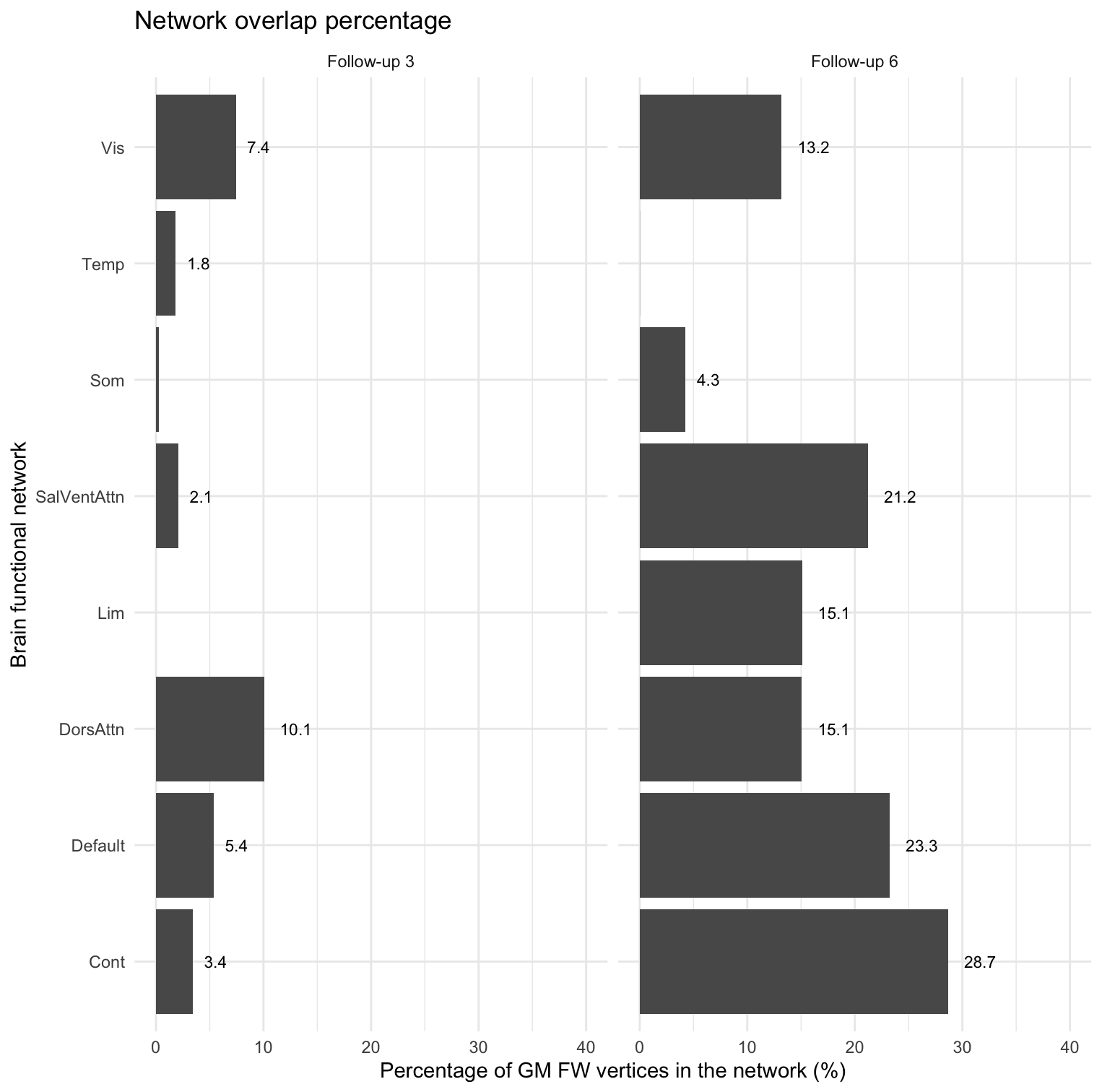

**Supplementary Figure 3.** Percentage of vertex overlap between major brain functional networks (Schaefer, 2018) and statistically significant grey matter free water (GM FW) clusters associated with higher DASH adherence in early elderhood (follow-up 3) and late life (follow-up 6).

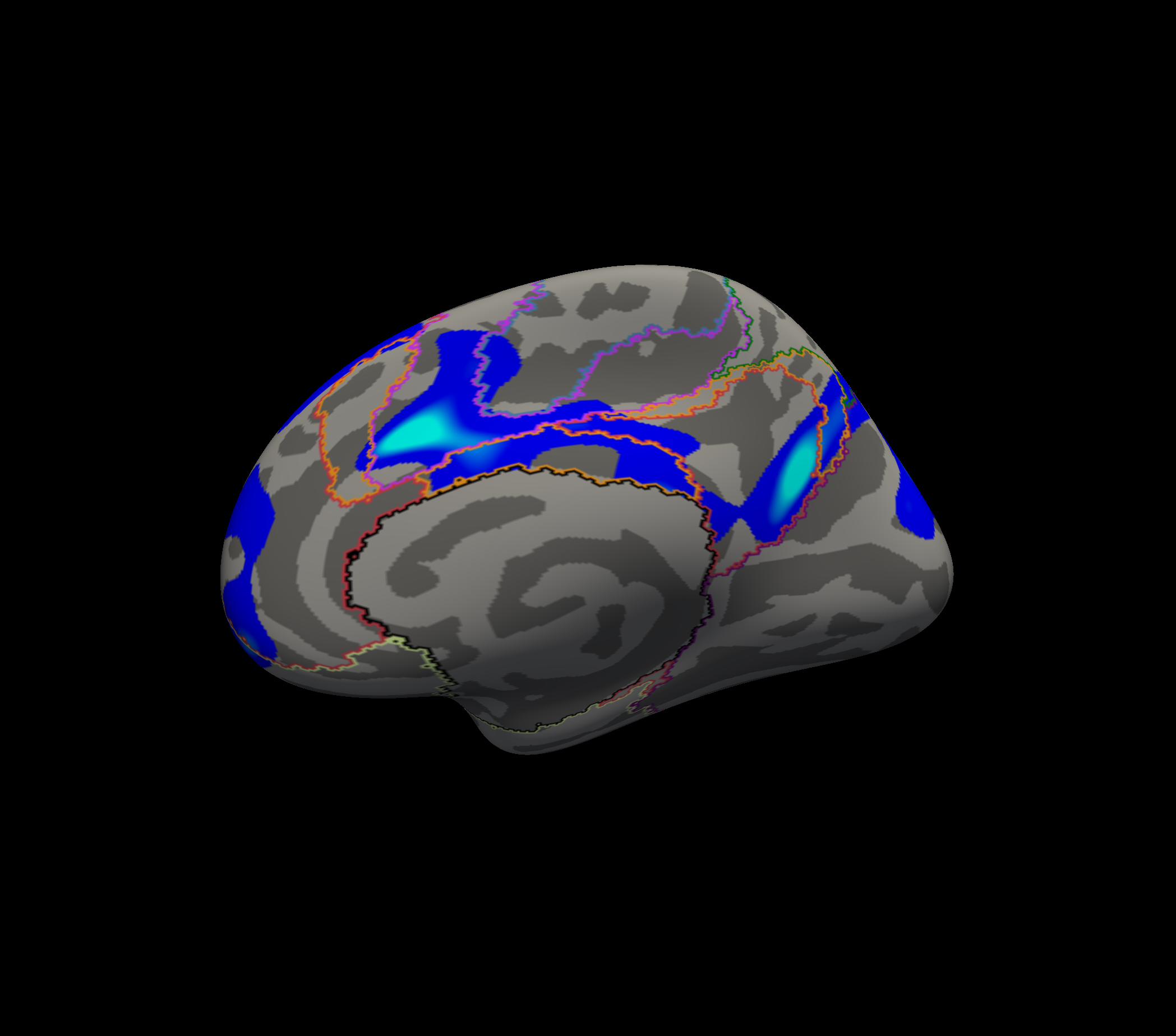

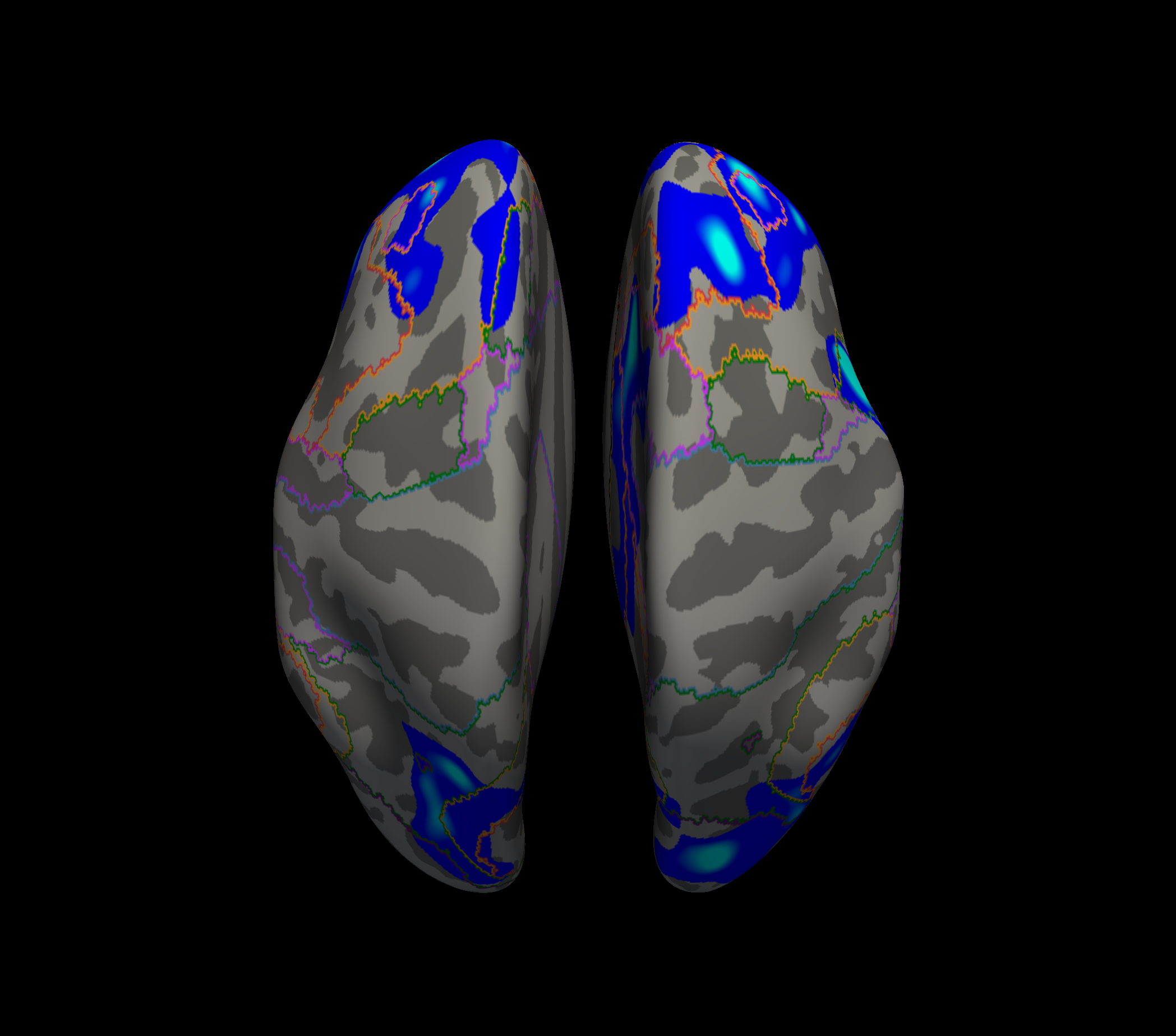

**Supplementary Figure 4.** Cross-sectional associations between higher late-life DASH adherence and lower late-life cortical grey matter free water. Colour represents p values of suprathreshold vertices (p = .001 to .05, brighter colour indicates smaller p value).

F. **Supplementary Results – Regional white matter hyperintensities**

Bullseye parcellation of WMH volume followed by factor analytic decomposition identified seven spatial components of regional WMHs (**Supplementary Fig. 5**). Among these components, WMHs in the white matter around deep basal ganglia and anterior periventricular zones were significantly associated with DASH adherence in early elderhood in the discovery Model 1 (**Supplementary Table 9**). The association with deep basal ganglia WMHs remained significant in models after additional covariate adjustment, while the association with anterior periventricular WMHs was attenuated (**Supplementary Table 10**).

| *Supplementary Table 9. Association between early elderhood DASH adherence and regional white matter hyperintensities adjusting for basic demographic covariates (Model 1)* | | | | | | | |
| --- | --- | --- | --- | --- | --- | --- | --- |
| DASH | Spatial component | *b* | *SE* | *t* | *p* | p_fdr | 95% CI |
| Early elderhood | **Anterior peri-ventricular** | **-0.04** | **0.02** | **-2.50** | **.012*** | **.044*** | **[-0.08, -0.01]** |
|  | **Deep basal ganglia** | **-0.02** | **0.01** | **-3.11** | **.002**** | **.013*** | **[-0.04, -0.01]** |
|  | Frontoparietal | -0.02 | 0.02 | -1.09 | .275 | .405 | [-0.05, 0.01] |
|  | Left horn | -0.01 | 0.01 | -1.08 | .281 | .405 | [-0.04, 0.01] |
|  | Temporal | -0.02 | 0.02 | -1.04 | .298 | .405 | [-0.06, 0.02] |
|  | Parietal | -0.02 | 0.02 | -0.94 | .347 | .405 | [-0.05, 0.02] |
|  | Right horn | -0.01 | 0.01 | -0.70 | .481 | .481 | [-0.03, 0.01] |
| *Note:* Model 1 controlled for total food energy measured in baseline, age at follow-up 6, handedness, sex, dialect group, levels of education, and intracranial volume. b = beta coefficient, SE = standard error, CI = 95% confidence intervals, p_fdr = multiple comparisons adjusted for 7 WMH spatial components. * p < .05, ** p < .01, *** p < .001 | | | | | | | |

| *Supplementary Table 10. Statistical summaries of the associations between early elderhood DASH and deep basal ganglia and anterior peri-ventricular WMH components* | | | | | | | |
| --- | --- | --- | --- | --- | --- | --- | --- |
| DASH | Component | Model | *b* | *SE* | *t* | *p* | 95% CI |
| Early elderhood | **Deep basal ganglia** | **2** | **-0.018** | **.007** | **-2.43** | **.016*** | **[-0.03, -0.001]** |
|  | **Deep basal ganglia** | **3** | **-0.016** | **.007** | **-2.18** | **.030*** | **[-0.03, -0.001]** |
|  | Anterior peri-ventricular | 2 | -0.035 | .018 | -1.90 | .058 | [-0.07, 0.001] |
|  | Anterior peri-ventricular | 3 | -0.031 | .019 | -1.65 | .099 | [-0.07, 0.01] |
| *Note:* Model 1 controlled for total food energy measured in baseline, age at follow-up 6, handedness, sex, dialect group, levels of education, and intracranial volume; Model 2 controlled for Model 1 and key covariates measured at the same time point as DASH. Model 3 controlled for Model 2 and key covariates measured at follow-up 6. * p < .05 | | | | | | | |

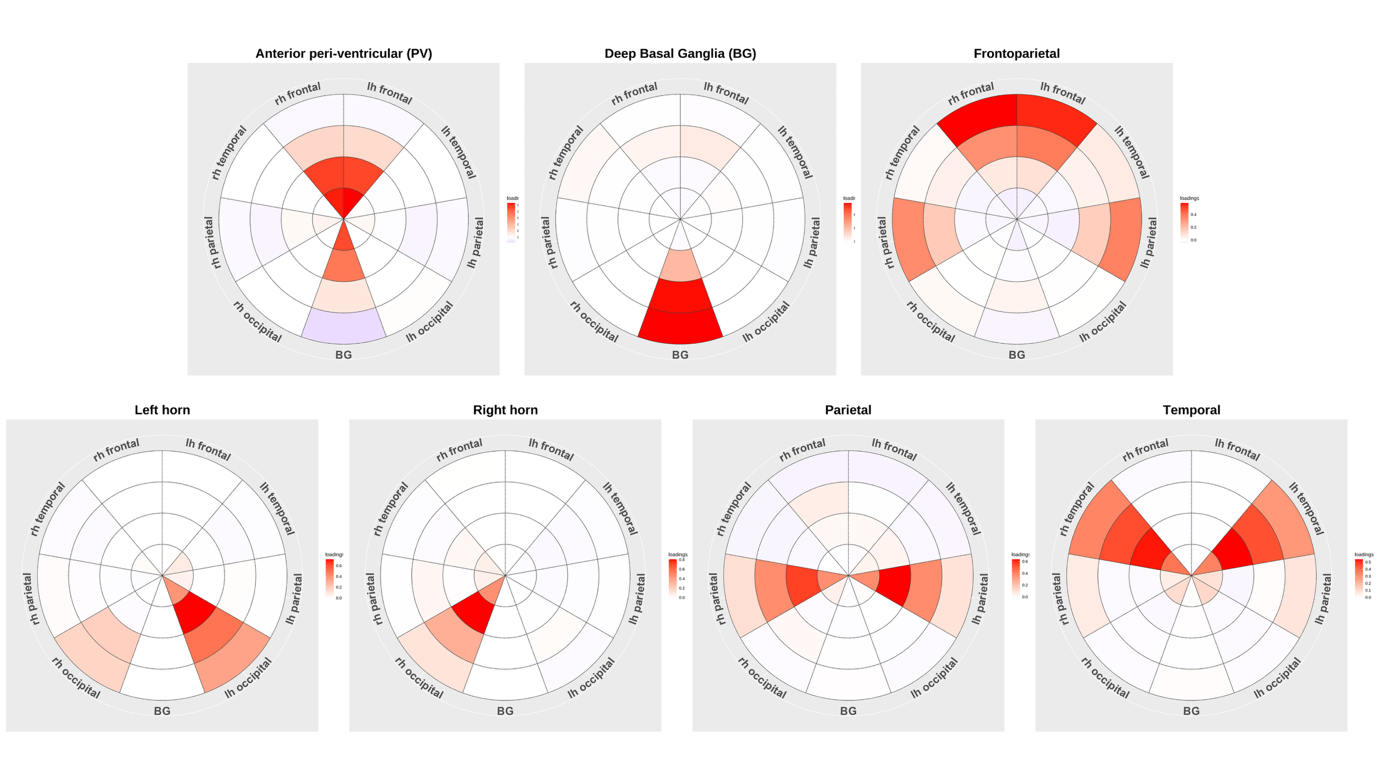
**Supplementary Figure 5.** Bullseye representation of the seven regional white matter hyperintensities (WMHs) components identified in the exploratory graph analysis (EGA). Concentric rings represent four equidistant zones between the ventricle (innermost) and the grey matter-white matter boundary (outmost), transversing from peri-ventricular to deep and juxta-cortical depths of the white matter. These zones are further divided according to lobes (frontal, temporal, parietal, occipital, and basal ganglia) and hemisphere (except basal ganglia). Stronger loadings (greater WMH burden) are indicated by warmer colours. Right hemisphere is displayed on the left. Abbreviations: lh = left hemisphere, rh = right hemisphere.

F. **Supplementary Results – Mediation analyses**

Across models with progressive covariate adjustment, both white matter volume and cortical grey matter free water were consistently associated with verbal memory in cross-sectional analyses at late life (**Supplementary Table 11**).

| *Supplementary Table 11. Associations between late life verbal memory and white matter volume (WMV) and cortical grey matter free water (cortical GM FW)* | | | | | | | |
| --- | --- | --- | --- | --- | --- | --- | --- |
| DASH | Measure | Model | *b* | *SE* | *t* | *p* | 95% CI |
| Midlife | WMV | 1 | 28.48 | 8.16 | 3.49 | .001*** | [12.46, 44.50] |
|  |  | 2 | 27.82 | 8.35 | 3.33 | .001*** | [11.42, 44.22] |
|  |  | 3 | 24.90 | 8.48 | 2.94 | .003** | [8.25, 41.54] |
| Early elderhood | Cortical  GM FW | 1 | -0.36 | .098 | -3.65 | < .001*** | [-0.55, -0.16] |
|  |  | 2 | -0.33 | 0.10 | -3.24 | .001** | [-0.54, -0.13] |
|  |  | 3 | -0.31 | 0.10 | -2.97 | .003** | [-0.52, -0.11] |
| *Note:* Model 1 controlled for total food energy measured in baseline, age at follow-up 6, handedness, sex, dialect group, levels of education, and intracranial volume; Model 2 controlled for Model 1 and key covariates measured at the same time point as DASH. Model 3 controlled for Model 2 and key covariates measured at follow-up 6. The appropriate DASH score was also included in each model. b = beta coefficient, SE = standard error, CI = 95% confidence intervals. *** p < .01 | | | | | | | |

G. **Supplementary Results - DASH trajectory analyses**

Across models specifying one to seven classes, latent class growth analysis identified six DASH trajectory groups as the optimal solution (**Supplementary Table 12**). These trajectories ranged from consistently low to consistently high DASH adherence, with two groups demonstrating improvement in their diet after midlife (from low to average adherence). Additional groups included a consistently average group with a slight post-midlife decline, and a group that began with low-to-average adherence in midlife but showed a marked decline thereafter (**Supplementary Table 13**).

To establish whether these trajectory groups differed meaningfully in cognitive and brain outcomes, we assessed group effects in multiple regression models for verbal memory, white matter volume (WMV), and cortical grey matter free water (GM FW) using omnibus F-tests. All three outcomes were significantly associated with trajectory group membership (verbal memory: F(5,814) = 2.81, p = .016; WMV: F(5,814) = 3.62, p = .003; cortical GM FW: F(5,804) = 2.77, p = .018). Our primary analyses focused on the potential cognitive and brain health benefits among the two improving trajectory groups in late life. Comprehensive summaries for all trajectory groups are provided for verbal memory (**Supplementary Table 14)**, white matter volume (**Supplementary Table 15**), and cortical grey matter free water (**Supplementary Table 16)**, including statistical significance adjusted for all pairwise comparisons with each anchoring group using FDR q = .05.

| *Supplementary Table 12. Latent class growth analysis (LCGA) model fit summary. Best values of each fit indices are bolded* | | | | | | | | | | | | |
| --- | --- | --- | --- | --- | --- | --- | --- | --- | --- | --- | --- | --- |
| *g* | loglik | AIC | BIC | saBIC | entropy | p(1) | p(2) | p(3) | p(4) | p(5) | p(6) | p(7) |
| 1 | -8,764 | 17,533 | 17,548.16 | 17,538.63 | 1.00 | 100 |  |  |  |  |  |  |
| 2 | -8,536 | 17,084 | 17,113.11 | 17,094.05 | 0.63 | 56 | 44 |  |  |  |  |  |
| 3 | -8,493 | 17,003 | **17,047.06** | 17,018.48 | 0.60 | 29 | 16 | 55 |  |  |  |  |
| 4 | -8,485 | 16,994 | 17,053.13 | 17,015.02 | 0.51 | 20 | 30 | 22 | 28 |  |  |  |
| 5 | -8,476 | 16,982 | 17,055.90 | 17,008.26 | 0.54 | 6 | 18 | 23 | 14 | 39 |  |  |
| 6 | -8,463 | **16,963** | *17,050.89* | **16,993.73** | 0.57 | 16 | 10 | 27 | 11 | 6 | 30 |  |
| 7 | -8,462 | 16,967 | 17,069.62 | 17,002.93 | 0.53 | 17 | 9 | 34 | 10 | 9 | 15 | 5 |
| *Note.* Abbreviations: g = Number of groups fitted, loglik = log likelihood, saBIC = sample size adjusted BIC, p(.) = percentage of participants in each group. Note that the percentages in p(.) are before labelling of the groups. | | | | | | | | | | | | |

| *Supplementary Table 13. The characteristics of the six DASH adherence trajectory groups* | | | | | | |
| --- | --- | --- | --- | --- | --- | --- |
| Class | *N* | Parameter | *b* | *SE* | *W* | *p* |
| 1 Consistently low | 49 | Intercept | 13.91 | 0.80 | 17.33 | < .001*** |
|  |  | Slope | 0.97 | 0.89 | 1.09 | .277 |
| 2 Low decreasing | 108 | Intercept | 19.71 | 0.62 | 31.87 | < .001*** |
|  |  | **Slope** | **-4.17** | **0.88** | **-4.76** | **< .001***** |
| 3 Low increasing | 225 | Intercept | 17.15 | 0.49 | 35.12 | < .001*** |
|  |  | **Slope** | **3.83** | **0.64** | **5.96** | **< .001***** |
| 4 Consistently average | 278 | Intercept | 22.79 | 0.54 | 42.11 | < .001*** |
|  |  | Slope | -1.20 | 0.66 | -1.83 | .067 |
| 5 Average increasing | 143 | Intercept | 21.43 | 0.68 | 31.41 | < .001*** |
|  |  | **Slope** | **4.59** | **0.85** | **5.38** | **< .001***** |
| 6 Consistently high | 89 | Intercept | 27.00 | 0.68 | 39.67 | < .001*** |
|  |  | Slope | -0.83 | 0.84 | -0.99 | .321 |
| *Note.* b = beta coefficient, SE = standard error, W = Wald’s test from hlme().* p < .05, ** p < .01, *** p < .001 | | | | | | |

| *Supplementary Table 14. Statistical summary of differences in late-life verbal memory between the best and worst DASH trajectory groups and the intermediate groups* | | | | | | |
| --- | --- | --- | --- | --- | --- | --- |
| Target trajectory group | Reference trajectory group | Estimate | SE | *t* | *p* | *p_fdr* |
| Low decreasing | Consistently high | -2.8 | 1.21 | -2.31 | .021* | .038* |
| **Low increasing** | Consistently high | -2.4 | 1.05 | -2.28 | .023* | .038* |
| Consistently average | Consistently high | -1.68 | 1 | -1.67 | .095 | .119 |
| **Average increasing** | Consistently high | -1.05 | 1.12 | -0.94 | .347 | .347 |
| Low decreasing | Consistently low | 2.39 | 1.41 | 1.7 | .09 | .09 |
| **Low increasing** | Consistently low | 2.78 | 1.31 | 2.13 | .034* | .042* |
| Consistently average | Consistently low | 3.51 | 1.31 | 2.67 | .008** | .013* |
| **Average increasing** | Consistently low | 4.14 | 1.42 | 2.92 | .004** | .009** |
| *Note*: The comparison between Consistently high and Consistently low is omitted. SE = standard error, p_fdr = multiple comparisons adjusted for 4 pairwise comparison with each anchor group. * p < .05, ** p < .01 | | | | | | |

| *Supplementary Table 15. Statistical summary of differences in late-life white matter volume between the best and worst DASH trajectory groups and the intermediate groups* | | | | | | |
| --- | --- | --- | --- | --- | --- | --- |
| Target trajectory group | Reference trajectory group | Estimate | SE | *t* | *p* | *p_fdr* |
| Low decreasing | Consistently high | -1.56 | 0.5 | -3.15 | .002** | .004** |
| **Low increasing** | Consistently high | -1.23 | 0.43 | -2.86 | .004** | .007** |
| Consistently average | Consistently high | -0.61 | 0.41 | -1.48 | .14 | .14 |
| **Average increasing** | Consistently high | -0.98 | 0.46 | -2.16 | .031* | .039* |
| Low decreasing | Consistently low | 0.51 | 0.58 | 0.89 | .376 | .376 |
| **Low increasing** | Consistently low | 0.85 | 0.54 | 1.57 | .118 | .147 |
| Consistently average | Consistently low | 1.47 | 0.54 | 2.72 | .007** | .017* |
| **Average increasing** | Consistently low | 1.09 | 0.58 | 1.88 | .061 | .101 |
| *Note*: Estimate and standard error (SE) are multiplied by 100 for visualisation only. The comparison between Consistently high and Consistently low is omitted. p_fdr = multiple comparisons adjusted for 4 pairwise comparison with each anchor group. * p < .05, ** p < .01 | | | | | | |

| *Supplementary Table 16. Statistical summary of differences in late-life cortical grey matter free water between the best and worst DASH trajectory groups and the intermediate groups* | | | | | | |
| --- | --- | --- | --- | --- | --- | --- |
| Target trajectory group | Reference trajectory group | Estimate | SE | *t* | *p* | *p_fdr* |
| Low decreasing | Consistently high | 12.77 | 4.32 | 2.96 | .003** | .013* |
| **Low increasing** | Consistently high | 9.05 | 3.74 | 2.42 | .016* | .020* |
| Consistently average | Consistently high | 8.49 | 3.53 | 2.41 | .016* | .020* |
| **Average increasing** | Consistently high | 3.63 | 3.92 | 0.93 | 0.355 | 0.355 |
| Low decreasing | Consistently low | -2.7 | 4.99 | -0.54 | .589 | .589 |
| **Low increasing** | Consistently low | -6.43 | 4.7 | -1.37 | .172 | .215 |
| Consistently average | Consistently low | -6.98 | 4.74 | -1.47 | .141 | .215 |
| **Average increasing** | Consistently low | -11.85 | 5.1 | -2.32 | .020* | .051 |
| *Note*: Estimate and standard error (SE) is multiplied by 1000 for visualisation only. The comparison between Consistently high and Consistently low is omitted. p_fdr = multiple comparisons adjusted for 4 pairwise comparison with each anchor group. * p < .05, ** p < .01 | | | | | | |

H. **Supplementary Results - Validation analyses**

Associations between DASH adherence and cognitive and brain measures were re-examined in a subset of healthier participants, excluding individuals with depressive symptoms or potential cognitive impairment (N = 104). Key findings, including those for verbal memory (**Supplementary Table 17**), white matter volume (**Supplementary Table 18**), deep basal ganglia WMH (**Supplementary Table 19**), cortical free water (**Supplementary Fig. 6**), and DASH trajectory comparisons with anchor groups (**Supplementary Table 20-22**) remained consistent. Mediation effects of white matter volume (indirect effect = .029, SE = .013, 95% CI = [.008, .057], 12.7% mediated) and cortical free water (indirect effect = .024, SE = .011, 95% CI = [.004, .049], 11.0% mediated) also remained consistent.

Grey matter morphometry showed the greatest variability, with thickness of the left precentral cortex, rather than the precuneus, showing a positive association with DASH adherence in early elderhood. The association with peri-ventricular WMH was also further attenuated.

| *Supplementary Table 17. Association between DASH adherence and cognitive performance in late life in a healthier subgroup, controlling for baseline and late life covariates (Model 3)* | | | | | | |
| --- | --- | --- | --- | --- | --- | --- |
| DASH | Cognitive domain | *b* | *SE* | *t* | *p* | 95% CI |
| Midlife | Verbal memory | 0.23 | 0.06 | 3.57 | .001** | [0.10, 0.36] |
| Early elderhood | Verbal memory | 0.21 | 0.07 | 3.25 | .001** | [0.08, 0.34] |
| Late life | Verbal memory | 0.26 | 0.07 | 3.90 | < .001*** | [0.13, 0.39] |
| *Note.* Model 1 controlled for total food energy measured in baseline, age at follow-up 6, handedness, sex, dialect group, levels of education, and intracranial volume; Model 2 controlled for Model 1 and key covariates measured at the same time point as DASH. Model 3 controlled for Model 2 and key covariates measured at follow-up 6. b = beta coefficient, SE = standard error, CI = 95% confidence intervals. ** p < .01, *** p < .001 | | | | | | |

| *Supplementary Table 18. Cluster peak statistics of voxel-wise analysis between midlife DASH adherence and late life white matter volume (WMV) in a healthier subgroup, controlling for midlife and late life covariates (Model 3)* | | | | | |
| --- | --- | --- | --- | --- | --- |
| DASH | Cluster | X | Y | *Z* | Peak Z value |
| Midlife | 1 | 38 | 45 | 3 | 3.95 |
|  | 2 | -32 | 18 | 15 | 3.82 |
|  | 3 | 34 | -40 | 36 | 3.61 |
| *Note:* Model 1 controlled for total food energy measured in baseline, age at follow-up 6, handedness, sex, dialect group, levels of education, and intracranial volume; Model 2 controlled for Model 1 and key covariates measured at the same time point as DASH. Model 3 controlled for Model 2 and key covariates measured at follow-up 6. | | | | | |

| *Supplementary Table 19. Statistical summaries for deep basal ganglia and anterior peri-ventricular WMH components in a healthier subgroup, controlling for early elderhood and late- life covariates (Model 3)* | | | | | | |
| --- | --- | --- | --- | --- | --- | --- |
| DASH | Component | *b* | *SE* | *t* | *p* | 95% CI |
| Early elderhood | Deep basal ganglia | -0.020 | 0.008 | -2.61 | .009** | [-0.03, -0.01] |
|  | Anterior peri-ventricular | -0.022 | 0.020 | -1.10 | .270 | [-0.06, 0.02] |
| *Note:* Model 1 controlled for total food energy measured in baseline, age at follow-up 6, handedness, sex, dialect group, levels of education, and intracranial volume; Model 2 controlled for Model 1 and key covariates measured at the same time point as DASH. Model 3 controlled for Model 2 and key covariates measured at follow-up 6. b = beta coefficient, SE = standard error, CI = 95% confidence intervals. ** p < .01. | | | | | | |

| *Supplementary Table 20. Statistical summary of differences in late-life verbal memory between selected DASH trajectory groups, restricted to a healthier subgroup* | | | | | |
| --- | --- | --- | --- | --- | --- |
| Target trajectory group | Reference trajectory group | Estimate | SE | *t* | *p* |
| Low decreasing | Consistently high | -3.46 | 1.29 | -2.49 | .009** |
| **Low increasing** | Consistently high | -2.45 | 1.10 | -2.13 | .033* |
| Consistently average | Consistently high | -1.58 | 1.05 | -1.55 | .15 |
| **Average increasing** | Consistently high | -0.59 | 1.16 | -0.74 | .62 |
| Low decreasing | Consistently low | 2.17 | 1.53 | 1.42 | .16 |
| **Low increasing** | Consistently low | 3.18 | 1.43 | 2.22 | .027* |
| Consistently average | Consistently low | 4.05 | 1.45 | 2.79 | .005** |
| **Average increasing** | Consistently low | 5.04 | 1.55 | 3.25 | .001** |
| *Note:* The comparison between Consistently high and Consistently low is omitted. * p < .05, ** p < .01 | | | | | |

| *Supplementary Table 21. Statistical summary of differences in late-life white matter volume between the best and worst DASH trajectory groups and the intermediate groups, restricted to a healthier subgroup* | | | | | |
| --- | --- | --- | --- | --- | --- |
| Target trajectory group | Reference trajectory group | Estimate | SE | *t* | *p* |
| Low decreasing | Consistently high | -1.53 | 0.57 | -2.80 | .005** |
| **Low increasing** | Consistently high | -1.09 | 0.47 | -2.32 | .021* |
| Consistently average | Consistently high | -0.53 | 0.44 | -1.20 | .23 |
| **Average increasing** | Consistently high | -0.92 | 0.49 | -1.88 | .06 |
| Low decreasing | Consistently low | 0.34 | 0.64 | 0.54 | .59 |
| **Low increasing** | Consistently low | 0.78 | 0.59 | 1.32 | .19 |
| Consistently average | Consistently low | 1.34 | 0.60 | 2.22 | .026* |
| **Average increasing** | Consistently low | 0.95 | 0.64 | 1.48 | .14 |
| *Note*: Estimate and standard error (SE) is multiplied by 100 for visualisation only. The comparison between Consistently high and Consistently low is omitted. * p < .05, ** p < .01 | | | | | |

| *Supplementary Table 22. Statistical summary of differences in late-life cortical grey matter free water between the best and worst DASH trajectory groups and the intermediate groups, restricted to a healthier subgroup* | | | | | |
| --- | --- | --- | --- | --- | --- |
| Target trajectory group | Reference trajectory group | Estimate | SE | *t* | *p* |
| Low decreasing | Consistently high | 13.82 | 4.48 | 3.09 | .007** |
| **Low increasing** | Consistently high | 8.37 | 3.85 | 2.18 | .030* |
| Consistently average | Consistently high | 9.07 | 3.63 | 2.50 | .013* |
| **Average increasing** | Consistently high | 2.58 | 4.02 | 0.64 | .52 |
| Low decreasing | Consistently low | -1.91. | 5.22 | -0.36 | .72 |
| **Low increasing** | Consistently low | -7.35 | 4.90 | -1.50 | .13 |
| Consistently average | Consistently low | -6.55 | 4.97 | -1.34 | .18 |
| **Average increasing** | Consistently low | -13.14 | 5.32 | -2.47 | .014* |
| *Note*: Estimate and standard error (SE) is multiplied by 1000 for visualisation only. The comparison between Consistently high and Consistently low is omitted. * p < .05, ** p < .01 | | | | | |

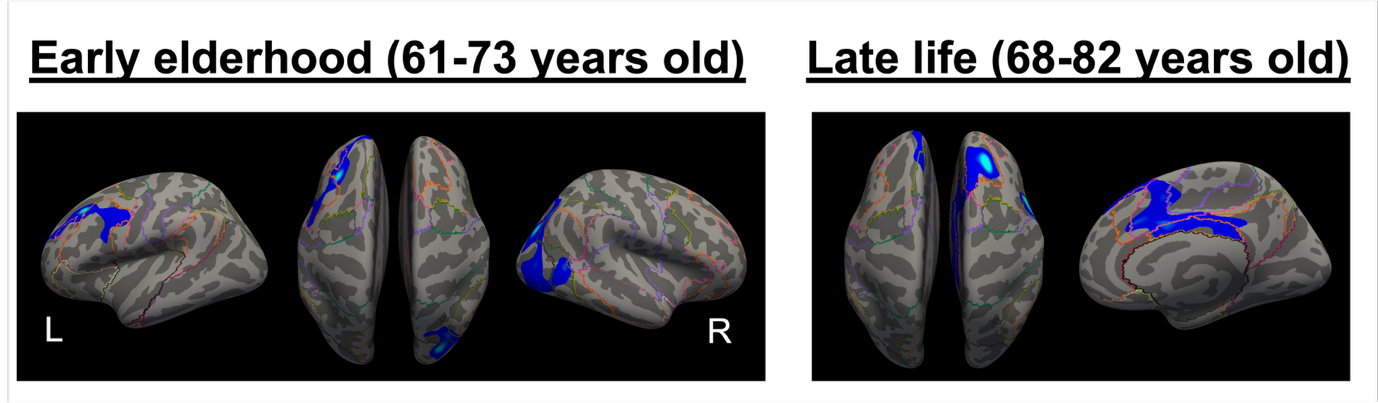

**Supplementary Figure 6.** Associations between higher life-stage DASH adherence and lower late-life cortical grey matter free water in a healthier subgroup. Colour represents p values of suprathreshold vertices (p = .001 to .05; brighter colour indicates smaller p value).
